# 3-Dimensional Organization and Dynamics of the Microsporidian Polar Tube Invasion Machinery

**DOI:** 10.1101/2020.04.03.024240

**Authors:** Pattana Jaroenlak, Michael Cammer, Alina Davydov, Joseph Sall, Mahrukh Usmani, Feng-Xia Liang, Damian C. Ekiert, Gira Bhabha

## Abstract

Microsporidia, a divergent group of single-celled eukaryotic parasites, harness a specialized harpoon-like invasion apparatus called the polar tube (PT) to gain entry into host cells. The PT is tightly coiled within the transmissible extracellular spore, and is about 20 times the length of the spore. Once triggered, the PT is rapidly ejected and is thought to penetrate the host cell, acting as a conduit for the transfer of infectious cargo into the host. The organization of this specialized infection apparatus in the spore, how it is deployed, and how the nucleus and other large cargo are transported through the narrow PT are not well understood. Here we use serial block-face scanning electron microscopy to reveal the 3-dimensional architecture of the PT and its relative spatial orientation to other organelles within the spore. Using high-speed optical microscopy, we also capture and quantify the entire PT germination process *in vitro*. Our results show that the emerging PT experiences very high accelerating forces to reach velocities exceeding 300 μm.s^-1^, and that firing kinetics differ markedly between species. Live-cell imaging reveals that the nucleus, which is approximately 7 times larger than the diameter of the PT, undergoes extreme deformation to fit through the narrow tube, and moves at speeds comparable to PT extension. Our study sheds new light on the 3-dimensional organization, dynamics, and mechanism of PT extrusion, and shows how infectious cargo moves through the tube to initiate infection.

---

Intracellular pathogens use a diverse array of mechanisms to enter and infect new host cells^1–3^. Microsporidia are a group of single-celled intracellular parasites that have developed one of the most dramatic, yet poorly understood, mechanisms of host cell invasion. Microsporidia are highly diverged from other extant eukaryotes, and are thought to be an early branching sister group to fungi^4,5^. Microsporidia infect a wide range of hosts, including nematodes^6,7^, insects^8,9^, and vertebrates^10,11^, including humans^12^. As obligate intracellular parasites with a reduced genome^13^, they are dependent on the host for replication^14,15^. Prior to exiting the host cell, microsporidia form spores, which are the only form of the organism that can survive outside of a host.

To initiate infection of a new cell, these parasites have evolved a unique, harpoon-like invasion apparatus called the polar tube (PT)^15–17^ that is present in all microsporidian species. The PT is tightly coiled within the dormant spore, resembling a spring^18,19^. When triggered, the PT transitions within a few hundred milliseconds from a spring-like coil to an extended linear tube, which in some species is more than 100 μm long^20^. The extended tube may penetrate or latch onto the target cell membrane to anchor the spore to the host cell^18,19,21,22^. The hollow PT is then poised to serve as a conduit for infectious material (also called sporoplasm) to enter the host cell and establish a replicative niche^23–25^. The entire process, from initiation of PT firing to the completion of cargo transport through the tube, is called spore germination.

The germination of microsporidian spores has fascinated biologists for more than 100 years^16^, yet many fundamental questions remain largely unexplored. While 2D TEM sections have revealed that parts of the PT are coiled in microsporidian spores^17,18,26^, the question of how the entire PT is configured within the spore, and its relation to other organelles in 3D is not well understood. The mechanisms that underlie the reconfiguration from a coil to a linear tube during PT extrusion are also a subject of considerable debate, yet this process has been challenging to study due to the stochastic firing of individual spores and the very fast (millisecond) timescales on which the germination process occurs^20,27^. Finally, the nucleus and other organelles of the microsporidian spore are thought to be translocated through the PT into the host cell. However, the PT is extremely narrow, raising the question: how does the infectious sporoplasm, 2-3 μm in size, move through a tube that is only 100 nm in diameter? Here we use a combination of serial block-face scanning electron microscopy (SBFSEM) and live-cell imaging to reveal the packaging of the long PT inside the much smaller spore, and its spatial relation to other organelles prior to PT germination. Furthermore, we unveil the dynamics of PT extrusion, and the mechanism of nuclear transport through the PT.

## Results

### 3D reconstruction of *A. algerae* spores

2D transmission electron microscopy (TEM) images of microsporidia have shown that the PT is arranged as a coil within the spore, alongside other organelles^18,26,28,29^. How the coils of the PT are organized in 3D and its spatial relationship to other organelles within the spore is less well understood. We used SBFSEM^30^ to generate 3D reconstructions of the PT in intact dormant spores of the microsporidian species *Anncaliia algerae*, which can infect both invertebrate and vertebrate hosts including humans^31– 34^. SBFSEM allows us to automate the sectioning and imaging of a series of 50 nm slices through a block of spores in a high throughput manner^35^. A typical field of view contained approximately 50 spores in several different orientations, from which we reconstructed 20 spores in 3D (**Fig. 1a**).

**Fig. 1.**
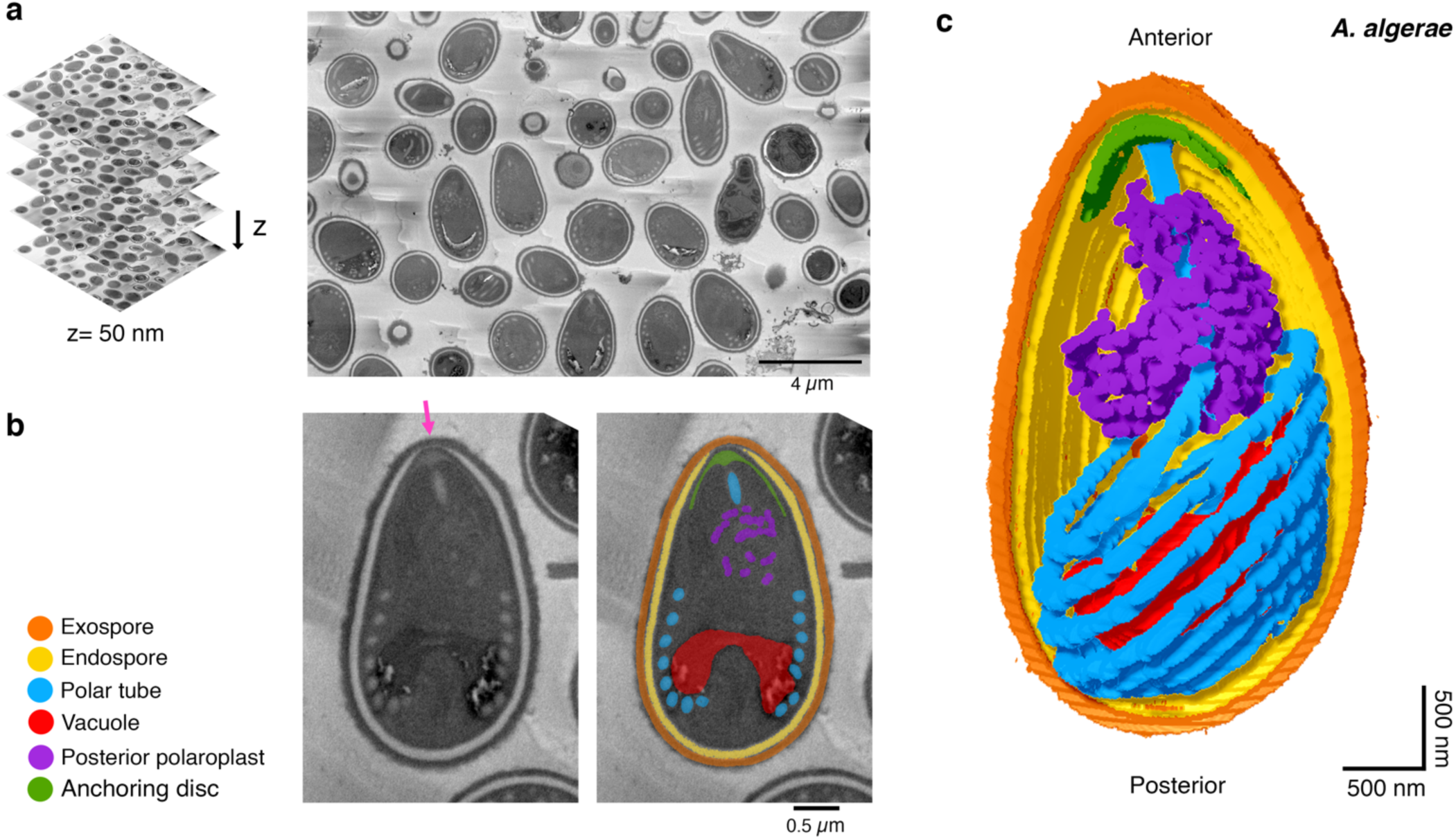
Serial block-face scanning electron microscopy imaging of intact *A. algerae* spores. **(a)** Representative data from SBFSEM imaging. Samples were serially sliced at 50 nm thickness (left), and images for a representative slice is shown (right). **(b)** Representative SBFSEM slice highlighting segmented organelles. Original micrograph is shown (left), as well as the same image with color overlays indicating segmented organelles (right): exospore (orange), endospore (yellow), PT (blue), vacuole (red), posterior polaroplast (purple), and anchoring disc (green). Magenta arrow indicates the thinnest part of the endospore layer where the anchoring disc is localized. **(c)** Representative 3D reconstruction of an *A. algerae* spore from SBFSEM data. Each color represents an individual organelle; color code as in **(b)**.

We segmented the outermost layer of the spore, the exospore, as well as the underlying chitinous endospore, which is adjacent to the plasma membrane (**Fig. 1b, Supplementary Video 1**). In the egg-shaped *A. algerae* spores, the exospore layer is uniformly 0.16 ± 0.03 μm thick, and the thickness of the endospore varies from 0.14 μm to < 0.08 μm towards the anterior tip of the spore. We found that the apical part of the endospore is the thinnest, as previously reported^28,36^, and this region has been hypothesized to be the site of initiation for PT firing^19,37^ (**Fig. 1b,c**). The spores are 3.9 ± 0.4 μm in length along the anterior-posterior (A-P) axis (**Supplementary Fig. 1a**), consistent with previous work^33^, and the average spore volume is 8.8 ± 1.4 μm^3^ (**Supplementary Fig. 1b**).

### Spatial organization of the polar tube in dormant *A. algerae* spores

We segmented the entire *A. algerae* PT to obtain spatial information on how it is organized in the dormant spore (**Fig. 1c, Supplementary Video 1**). The PT occupies only ∼3.5% of the total spore volume (**Supplementary Fig. 1b**), yet visually is the most striking feature of the spores when imaged by either SBFSEM or 2D TEM. As previously reported^38^, the PT can be divided into two main parts: 1) a linear segment that emanates from the anterior tip of the spore and extends towards the posterior end, and 2) a coil of PT around the middle and posterior end of the spore (**Fig. 2a**). At least two models for the connection between these parts have been proposed (**Supplementary Fig. 2**). In the first model, the linear segment extends nearly the length of the spore, and connects to the posterior end of the coiled region^39^. In the second model, the linear segment is shorter, and connects with the anterior end of the coiled region^37^. Our data show that the straight segment connects to the anterior end of the coiled region (**Fig. 2a**), consistent with the second model. The coils of the PT are approximately parallel to each other, but the stack of coils is tilted relative to the A-P axis of the spore (**Fig. 2b**). Here we found that the coiled segment of the PT consists of 7 turns on average (**Supplementary Fig. 1c**), in contrast to previous data which suggested 8-11 coils^33^, perhaps due to differences in the source or propagation of the spores.

**Fig. 2.**
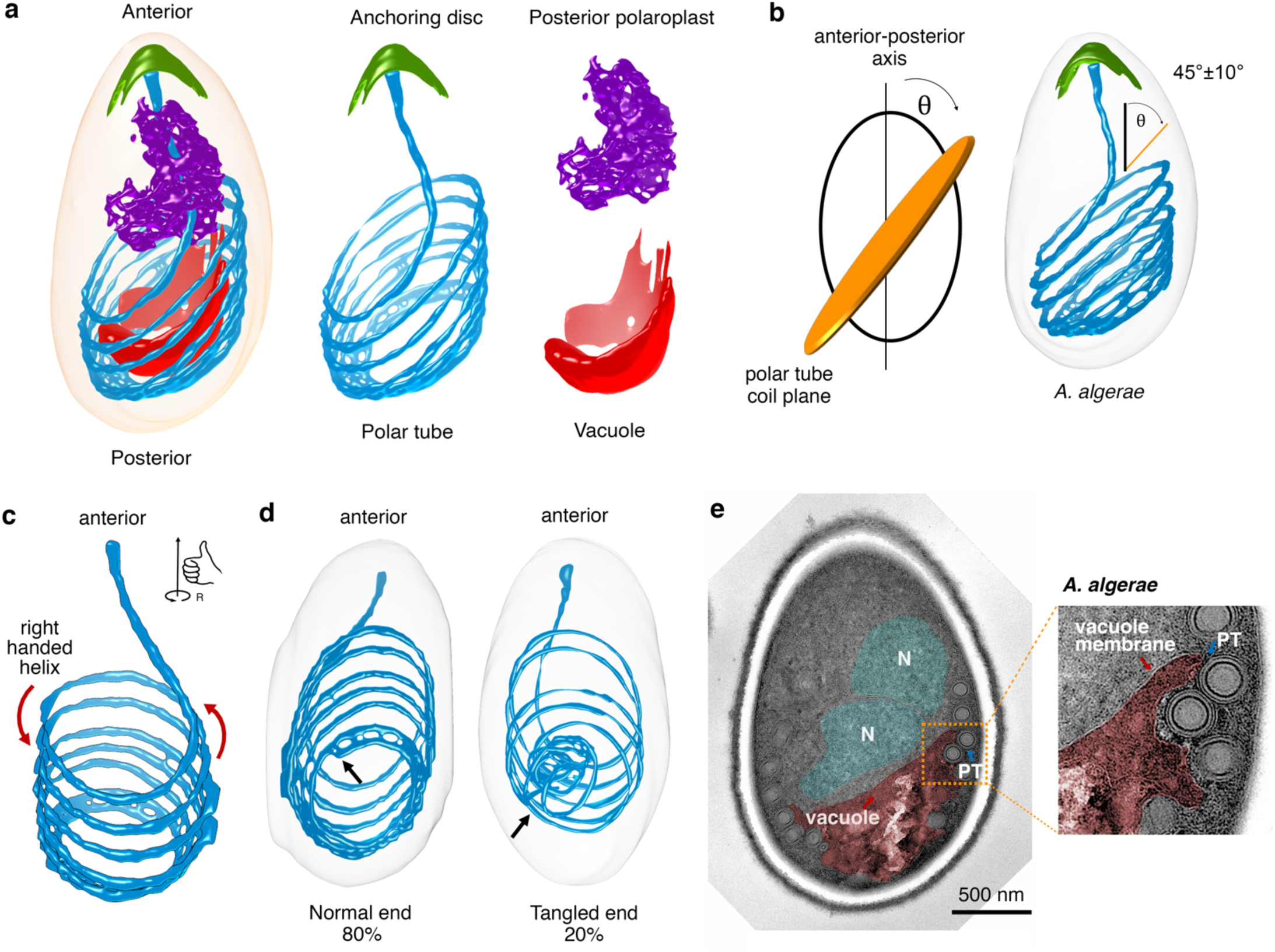
Configuration of the PT and other organelles in intact *A. algerae* spores. **(a)** Representative 3D reconstruction of an *A. algerae* spore showing the relative orientations of the PT (blue), anchoring disc (green), posterior polaroplast (purple), and vacuole (red). **(b)** measurement of the angle of PT coils relative to the anterior-posterior (A-P) axis of the spore. (left) Schematic showing how the angle was measured between the A-P axis (black) and the PT coil plane (orange). (right) Representative spore with the average angle and standard deviation annotated (n=18). **(c)** Chirality of the *A. algerae* PT. Red arrows indicate the right-handed helix of the PT. **(d)** Heterogeneity of PT ends observed in *A. algerae* PTs (n = 20). Black arrows indicate the location of the PT end. **(e)** Transmission electron microscopy (TEM) section of an *A. algerae* spore showing interdigitation between vacuole membrane (red) and the PT, and position of the nuclei (N, cyan) relative to the vacuole. Inset shows the region boxed in orange.

Remarkably, in every spore examined the PT takes the form of a right-handed helix (**Fig. 2c**). In the absence of any mechanism to bias PT assembly, the expectation would be a 50:50 mixture of left-handed and right-handed helices. This strong bias towards a right-handed helix implies the presence of an organizing principle. One explanation is that handedness could arise from the physical properties of a polymer (possibly formed by the polar tube proteins^40–42^) acting as a key structural element of the PT. This would be consistent with other biopolymers — such as DNA, actin filaments, and microtubules — which adopt superhelical coils with characteristic twist and handedness under strain. Alternatively, this right-handed bias may arise from asymmetry in the machinery involved in the PT assembly process.

We observed heterogeneity at the posterior end of the PT among spores, leading to two main classes of PT ends (**Fig. 2d**). In the majority of spores, the PT ends abruptly with a clean, blunt posterior end, remaining well-aligned with the preceding coils. However, in a few spores we observed irregular, tangled ends (**Fig. 2d, Supplementary Fig. 3**). The significance of blunt versus tangled PT ends is unclear, but the tangled ends may, for example, result from abnormal development of the PT during spore formation^43^. It seems likely that these tangled ends pose a problem for PT extrusion, and if not resolved may result in incomplete germination and failed infection.

### Configuration of organelles surrounding the PT in *A. algerae*

In addition to the PT, we segmented other organelles, including the anchoring disc, vacuole, and posterior polaroplast (**Fig. 1b,c, Supplementary Video 1**). First, the anchoring disc forms an umbrella-shaped structure at the anterior tip of the spore underlying the thinnest region of the endospore, as previously described^28,44^ (**Fig. 1b,c**). The anchoring disc is thought to be the site of spore germination, and thus some rearrangement or disruption of the anchoring disc is required to allow egress of the PT. Second, the vacuole at the posterior end of the spore occupies approximately 6.9% of the spore volume, and is roughly bowl-shaped (**Fig. 2a, Supplementary Fig. 1b**). The vacuole has previously been shown to expand during the germination process, and one possibility is that it plays an active role in facilitating PT extension and the translocation of spore contents through the PT^27,45^. Our 2D TEM sections show that the vacuole is surrounded by a single membrane (**Fig 2e**). The convex side is in close proximity to the posterior pole, and the concave side faces towards the anterior (**Fig. 2a,e**). While there is no clear cellular structure interacting with the concave surface of the vacuole in our SBFSEM images, examination of 2D TEM images reveals that the nucleus often rests against this depression in the vacuole (**Fig. 2e**). Interestingly, the vacuolar membrane is tightly interdigitated between PT coils (**Fig. 2e, Supplementary Fig. 4a**). Third, the polaroplast is a multilayer membranous organelle that is thought to be important during the germination process, perhaps providing membrane to accommodate the increased cell surface area during cargo transport through the PT^45^. Typically, the polaroplast consists of two regions with different morphologies, as observed by 2D TEM^25,26^: anterior polaroplast and posterior polaroplast (**Supplementary Fig. 4**). Reconstruction of the posterior polaroplast showed that it snugly surrounds the linear part of the PT (**Fig. 2a**), possibly stabilizing this region and is well-positioned to contribute membrane to the extending tube as it exits the spore.

### Comparison of *E. hellem* and *A. algerae* 3D reconstructions

To assess whether the PT configuration and the relative orientation of other organelles are conserved in other microsporidian species, we carried out SBFSEM analysis of another human-infecting microsporidian species, *Encephalitozoon hellem* (**Fig. 3a**). We segmented the spore wall layers, PT, anchoring disc, and anterior polaroplast (**Fig. 3b, Supplementary Video 2**). 3D reconstructions showed that while the overall organization of organelles in *E. hellem* spores is similar to the organization in *A. algerae* spores, there are also some notable differences. First, in contrast to the egg-shaped *A. algerae* spores, the *E. hellem* spores are more cylindrical (**Fig. 3c**). The length of the *E. hellem* spores is 2.8 ± 0.3 μm along the A-P axis, and the volume is approximately half that of *A. algerae* (**Supplementary Fig. 1a,b**). Second, the spacing between PT coils is smaller in *E. hellem* (an average distance of 0.12 ± 0.03 μm, compared with 0.22 ± 0.02 μm for *A. algerae* (**Fig. 3d**)), resulting in a more tightly packed coil. Third, in most *E. hellem* spores, the anchoring disc is located off-center with respect to the apical tip of the spore, rather than at the center of the apical tip as in *A. algerae* (**Fig. 3e**). The region of the spore wall surrounding the anchoring disc is the thinnest part (**Fig. 1b,3b**), regardless of whether it is at the apical tip, or off-centered, and is where the PT is expected to exit the spore. Overall, the SBFSEM results provide insights into how the PT is packed inside the spore, and these data provide a static snapshot of the PT in its pre-germination state.

**Fig. 3.**
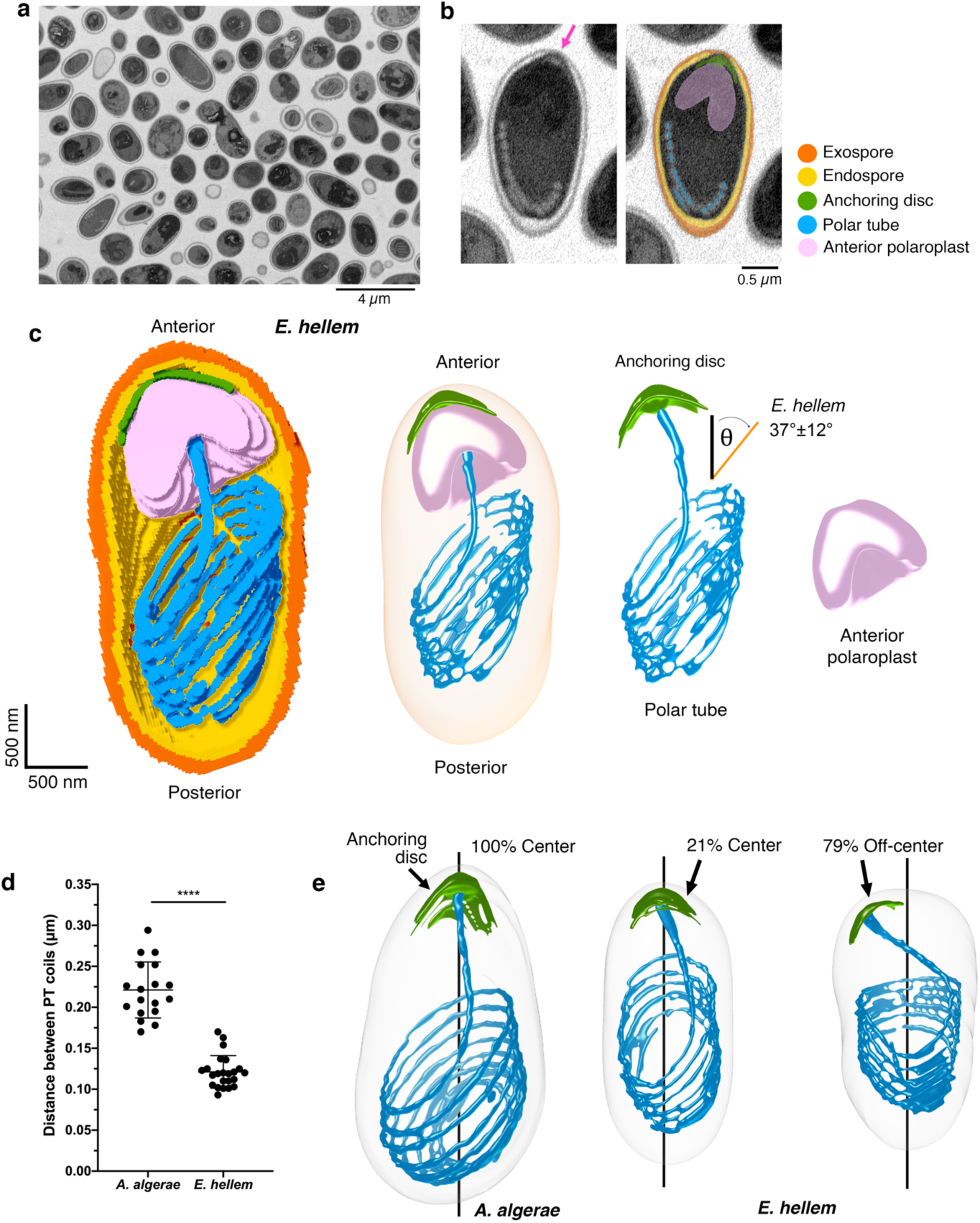
Serial block-face scanning electron microscopy imaging of *E. hellem* spores and comparison with *A. algerae*. **(a)** A representative slice from SBFSEM imaging of *E. hellem* spores. **(b)** Representative SBFSEM slice highlighting segmented organelles. Original micrograph is shown (left), as well as the same image with color overlays indicating segmented organelles (right): exospore (orange), endospore (yellow), PT (blue), vacuole (red), anterior polaroplast (pink), and anchoring disc (green). Magenta arrow indicates the thinnest part of the endospore layer where the anchoring disc is localized. **(c)** Representative 3D reconstruction of an *E. hellem* spore from SBFSEM data. Each color represents an individual organelle; color code as in (b). The average angle of *E. hellem* PT coils relative to the A-P axis is annotated with a standard deviation (n=20). The angle measurement was performed as in **Fig. 2b. (d)** Quantification of the distance between the PT coils in *A. algerae* and *E. hellem*. Error bars represent standard deviation (n=18 for *A. algerae* and n=20 for *E. hellem*), ****p<0.0001 (unpaired Student’s t-test). **(e)** Position of the anchoring disc (AD, green) relative to the spore coat. Black line indicates the anterior-posterior (A-P) axis of the spore. Coincidence of the AD and the A-P axis was scored as ‘center’, indicating that the AD is at the apical tip of the spore. Separation of the AD from the A-P axis was scored as “off-center”, indicating that the AD is not centered at the apical tip of the spore (n=18 for *A. algerae* and n=20 for *E. hellem*).

### Kinetics of polar tube germination

In order to infect the host, microsporidia must first extend the PT to penetrate a target cell. This process occurs extremely rapidly^20,27^, making it challenging to image in real time. To better understand the kinetics and mechanics of PT germination, we performed high-speed, live-cell imaging to capture *in vitro* PT germination events in three microsporidian species that infect humans: *A. algerae, E. hellem*, and *Encephalitozoon intestinalis*. Although the *in vivo* triggers for PT firing are not well understood, *in vitro* triggers have been reported^46–49^. We used small variations of these conditions to optimize PT firing *in vitro* for our light microscopy assay (**see Methods**), and captured the entire germination process, including release of the cargo after transport through the PT (**Fig. 4a, Supplementary Videos 3, 4, 5**). In all three species, we observed three distinct phases of the germination process: 1) PT elongation, 2) a static phase, where the PT length does not change, and 3) emergence of cargo at the distal end of the PT. PT elongation occurs rapidly (**Fig. 4b, Supplementary Fig. 5,6,7**); the time to reach 90% PT extension (T_EXT90_) is significantly shorter in *E. intestinalis* (T_EXT90_ 160 ± 20 ms; **Fig. 4c**) and *E. hellem* (T_EXT90_ 290 ± 200 ms) compared with *A. algerae* (T_EXT90_ 830 ± 170 ms). The entire germination process, from the start of PT extrusion to cargo ejection, is completed in less than 500 ms in *E. hellem* and *E. intestinalis*, and in approximately 1.6 seconds in *A. algerae*. We observed that the PT frequently emerges from the center of the apical region in *A. algerae* and off-center in *E. hellem* and *E. intestinalis* (**Supplementary Fig. 8**). This is consistent with the preferential positioning of the anchoring disc in our SBFSEM data, at the center of the apical tip in *A. algerae* and off-center in *E. hellem* spores, consistent with the idea that the anchoring disc position determines the PT exit site.

**Fig. 4.**
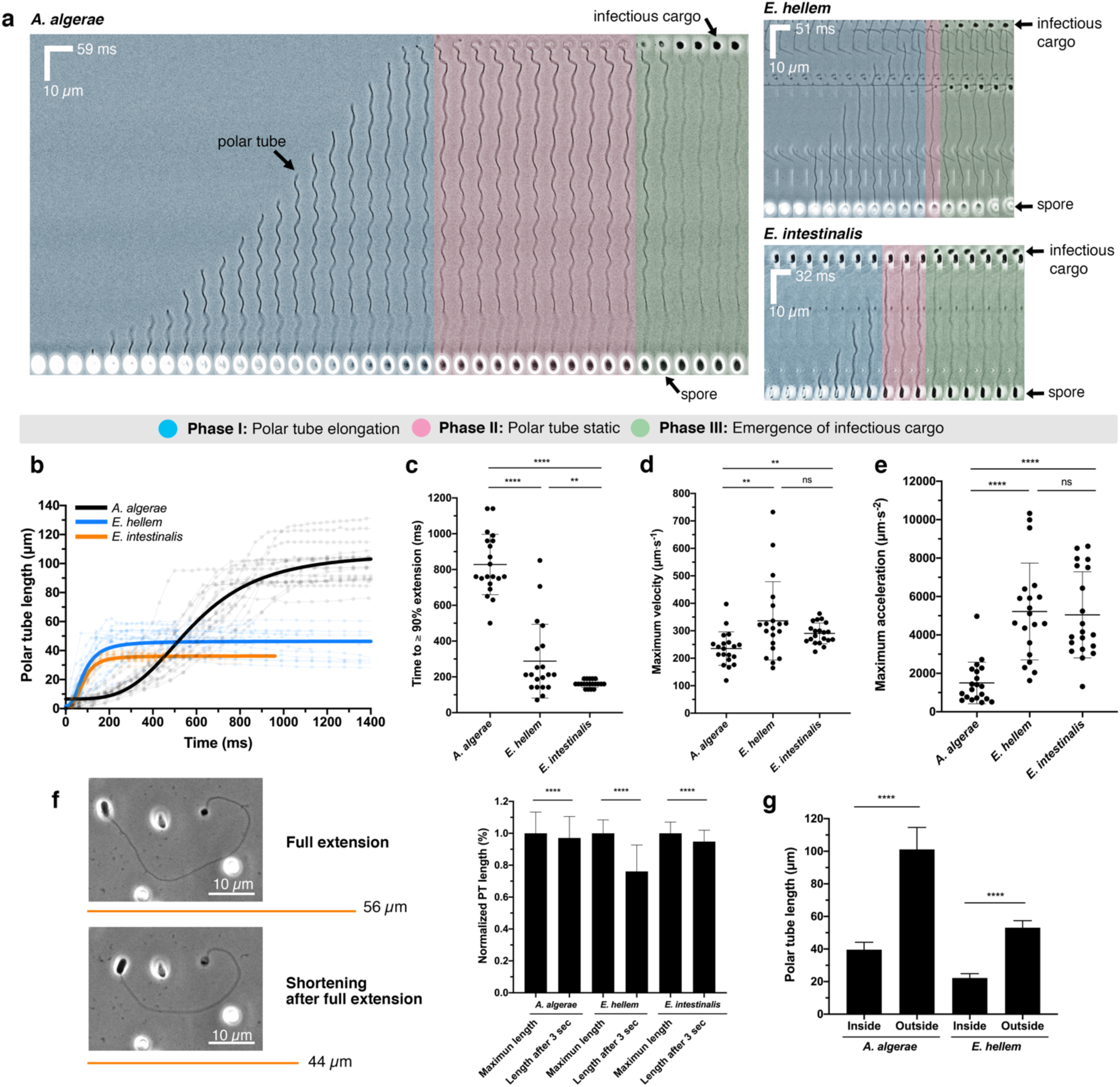
Live-cell imaging of PT germination. **(a)** Kymographs showing representative spore germination events from three microsporidia species: *A. algerae, E. hellem*, and *E. intestinalis*. Scale bars for distance and time intervals are shown on each kymograph. Three phases of the germination process are color-coded. The *A. algerae* PT appears to have a wavy pattern as it emerges from the spore, possibly forming a spiral, while the *E. hellem* and *E. intestinalis* PTs are less wavy in nature. As infectious cargo emerges from the distal end of the PT, the amplitude of the waveform is reduced, as has been previously noted^20^. After the *A. algerae* PT is fully extended, the distal end of the fired PT is curved and forms a hook-like structure, as previously described^54^. See also **Supplementary Videos 3, 4, 5. (b)** Quantification of the PT length as a function of time for *A. algerae* (black), *E. hellem* (blue), and *E. intestinalis* (orange) (n=20 for each species). The raw data are shown as faint lines with time points indicated as circles. The overall trends of each species were fitted (**see Methods**) and are represented as thick lines in the corresponding color. **(c)** Time taken for PT to extend to ≥ 90% (T_EXT90_) of its maximum length (n=20 for each species). ****p<0.0001, **p=0.0096 (unpaired Student’s t-test). **(d)** Average maximum velocity of PT extension calculated as described in **Methods** (n=20 for each species). **p=0.006; ns, not significant (unpaired Student’s t-test). **(e)** Average maximum acceleration of PT extension (n=20 for each species). ****p<0.0001; ns, not significant (unpaired Student’s t-test). **(f)** Shortening of the *E. hellem* PT after the full extension (maximum length) has been reached and cargo ejected. (left) Representative phase-contrast micrographs are shown, and orange lines represent the length of the PT quantified from the micrographs. (right) Quantification of the maximum PT length (full extension) and the length at 3 sec after the cargo is ejected. The graph shows PT length normalized to average maximum PT length (n=20 for each species, see also **Fig. S9a**). The PT shortens by 3%, 24%, and 5% for *A. algerae, E. hellem*, and *E. intestinalis*, respectively. ****p<0.0001 (paired Student’s t-test). **(g)** Comparison of the PT length in dormant spores, obtained from SBFSEM (labeled “inside”) and the maximum PT length after germination, obtained from live-cell optical microscopy experiments (labeled “outside”) (n=20 for each species, **see Methods** for how these measurements were made). ****p<0.0001 (unpaired Student’s t-test). All error bars in this figure represent SD.

In phase 1, the PT is fired and elongates to its maximum length (**Fig. 4b, Supplementary Fig. 5,6,7**). Our data show that on average, the maximum velocity (V_max_) of PT firing is 235 ± 61 μm.s^-1^ and the maximum acceleration (A_max_) is 1,503 ± 1,082 μm.s^-2^ in *A. algerae* (**Fig. 4d,e**). The V_max_ we observe is approximately double the previously reported value for this species^20^, which may reflect differences in *A. algerae* spores purified from different sources or more accurate measurements due to higher temporal resolution. In *E. hellem* and *E. intestinalis*, V_max_ and A_max_ are significantly higher than that measured for *A. algerae*. The V_max_ and A_max_ reaches 336 ± 142 μm.s^-1^ and 5,219 ± 2,521 μm.s^-2^ for *E. hellem*, and 290 ± 38 μm.s^-1^ and 5,045 ± 2,242 μm.s^-2^ for *E. intestinalis*, respectively (**Fig. 4d,e**). In contrast, other cellular processes involving movement are typically slower, such as organelle transport in cells by motor proteins (∼0.51 μm.s^-1^ for human kinesin^50^, ∼1 μm.s^-1^ for porcine dynein^51^), gliding motility of apicomplexan parasites (1-3 μm.s^-1^)^52^, and mobility of zoospores from chytrid fungi (∼104 μm.s^-1^)^53^.

In phase 2, the PT is fully extended and remains static at its maximum length: on average, 101, 53 and 36 μm for *A. algerae, E. hellem*, and *E. intestinalis* respectively (**Supplementary Fig. 9a**). In *A. algerae* this phase persists for 790 ± 360 ms (Supplementary Fig. 9b), and we hypothesize that during this time, the infectious cargo is traveling through the PT. If cargo transport begins once the tube is fully extended and all the species transport cargo at the same rate, we would expect phase 2 in *E. hellem* and *E. intestinalis* to be approximately 400 ms, since their PTs are approximately half the length of the *A. algerae* PT. However, we observe *E. hellem* and *E. intestinalis* spend significantly shorter time in this phase than predicted (60-140 ms; **Supplementary Fig. 9b**), suggesting that either cargo transport is faster in these species, or that cargo may begin moving through the PT before extension is completed. Intriguingly, for both *A. algerae* and *E. hellem*, we found that the fully extended PTs observed by light microscopy (LM) are more than twice the length of pre-germination PTs packaged inside the spore, as assessed by SBFSEM (**Fig. 4g, Supplementary Fig. 9c, see Methods**). Previous study has suggested that the PT is made of repetitive protein polymers^54^, and our observation raises the possibility that a significant conformational change may occur in some or all of these proteins between pre-germination and post-germination states, potentially leading to a change in PT length.

In phase 3, the cargo is expelled at the distal end of the PT, and appears as an approximately circular shape (**Fig. 4a**). The cargo remains attached to the tube for the duration of the experiment, and it is unclear what mediates this contact. There may be specific interactions between components of the PT and the cargo, or alternatively, some cargo may remain inside the tube itself, thereby creating a membranous bridge that leads to stable adhesion. When we continued imaging for several seconds after the cargo had been expelled, we observed that the *E. hellem* PT rapidly shortens (**Fig. 4f**). To assess this observation quantitatively, we measured the length of the PT after germination is complete (full PT extension), and also 3 s later. The PT of *E. hellem* shortens by 24%, while the length of the *A. algerae* and *E. intestinalis* PTs shorten by only 3% and 5% respectively (**Fig. 4f, Supplementary Fig. 9a, Supplementary Video 6**). This suggests that there are differences in the mechanics, plasticity and behavior of the PT even between closely related species such as *E. hellem* and *E. intestinalis*. Shortening of the *E. hellem* PT appears to be distributed across the length of the tube, as opposed to shortening due to retraction of the tube back inside the spore coat (**Supplementary Video 6**).

Occasionally, we observed incomplete germination of the PT in all three species, which we define as being stuck in phase 1 or phase 2: the PT may not be fully extended, or it is extended but no cargo is observed at the distal end of the tube (**Supplementary Fig. 8a,b, Supplementary Videos 7, 8, 9**). In these events, the time to maximum extension of the PT was longer than in productive germination events (**Supplementary Fig. 8c**). Incomplete PT firing events have been previously described^20^, and may represent spores that are not infectious, since cargo does not emerge at the distal end of the tube.

### Cargo transport through the tube

An enigma of the microsporidian invasion process is that the cargo to be transported through the tube is much larger than the diameter of the tube itself. While it is unclear precisely what is transferred from the spore to the host cell, at the very least the nucleus containing the parasite genome must be transported through the tube^26,55^. In the case of *A. algerae*, there are two nuclei^19^, and the diameter of each nucleus is about 0.7 μm, while the diameter of the PT is 100 nm^54^. This mismatch in scale must be overcome to achieve cargo transport through the tube. As tools for genetic modification and transgenesis are not yet available for microsporidia, labeling specific subcellular structures for live-cell imaging is challenging, and consequently tracking the movement of cargo through the polar tube has not been reported.

To track nuclear movement within the spore body during germination, we used NucBlue to stain the nucleus of intact spores, which yields sufficient signal for live-cell imaging (**Fig. 5a**). We used a modified version of our live-cell imaging assay, in which we pre-incubated dormant *A. algerae* spores with NucBlue, and tracked the nucleus during PT firing. We used dual detection of fluorescence and transmission to track the nucleus and to visualize the PT, respectively. Prior to nuclear exit from the spore, we clearly observed two distinct fluorescent lobes, corresponding to the two nuclei in *A. algerae* (**Fig. 5c, Supplementary Video 10**). The two nuclei briefly move around and rearrange inside the spore, perhaps in response to the rapid ejection of the PT, then enter the PT together, on a very fast timescale. Interestingly, while T_EXT90_ is 830 ms in *A. algerae* on average, nuclear translocation into the PT begins only ∼500 ms after the initiation of PT firing, suggesting that the tube is not fully extended prior to cargo transport (**Fig. 5c, Supplementary Fig. 10a**).

**Fig. 5.**
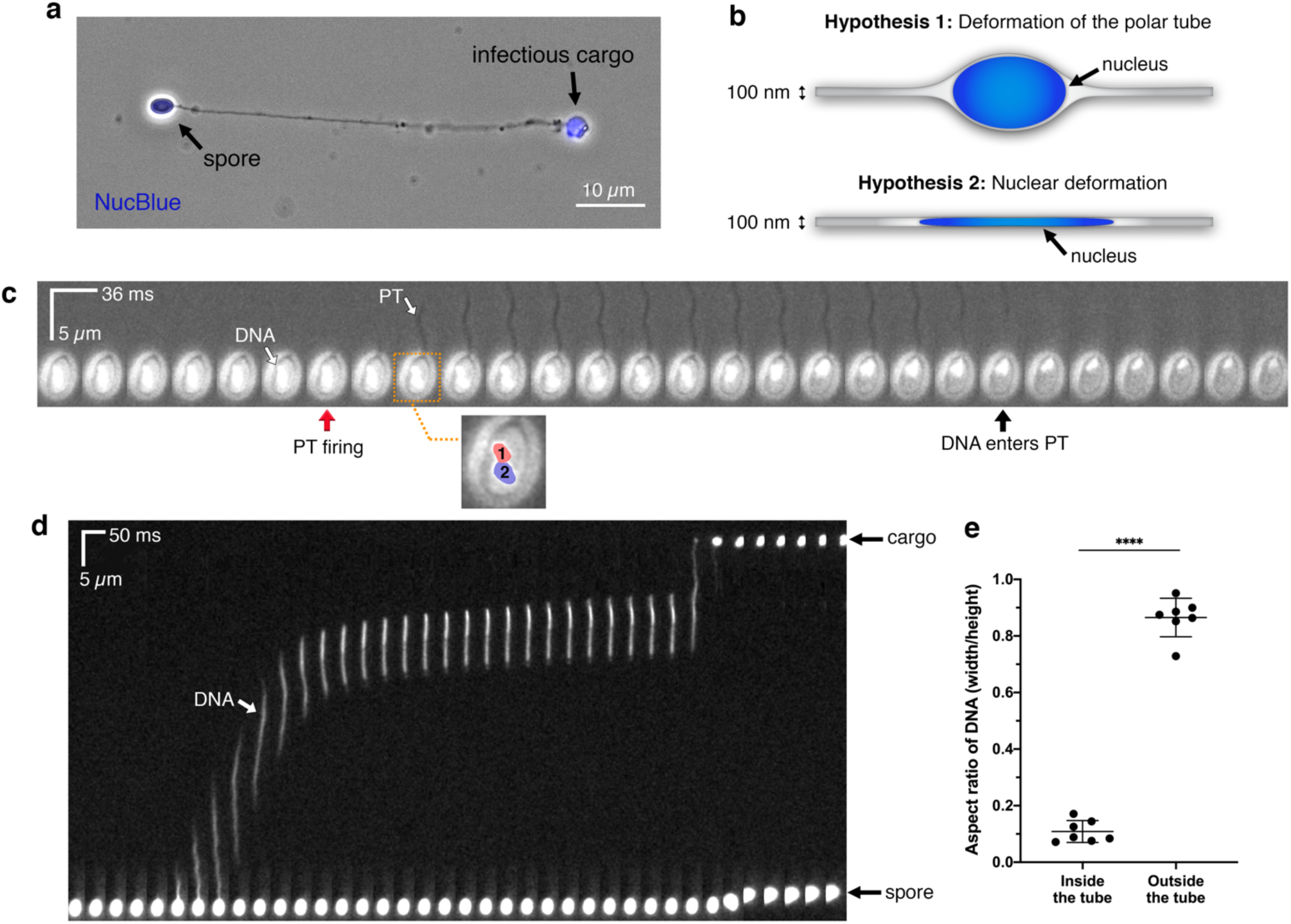
Live-cell imaging of nuclear transport through the PT. **(a)** Image of a fixed germinated *A. algerae* spore stained with a nuclear dye, NucBlue, overlaid with a phase-contrast image of the same spore. **(b)** Schematic diagrams show two possible hypotheses of how large nuclei may travel through the narrow PT. Nucleus is depicted in blue; in “hypothesis 1” it is drawn as an oval, to scale with the PT (nucleus diameter = ∼0.7 µm; PT diameter = ∼100 nm). **(c)** Time-lapse images of the two *A. algerae* nuclei inside the spore during PT germination, with a time interval of 36 ms. Nuclei are pre-stained with NucBlue, and white light was applied in order to observe the PT firing event simultaneously. Red arrow indicates the frame in which PT firing is initiated; black arrow indicates the frame in which the nuclei begin to leave the spore body. Inset highlights the two nuclei, labeled with numbers and color overlays, from the frame boxed in orange. **(d)** Kymograph of nuclear translocation through the PT, with a time interval of 50 ms. **(e)** Quantification of the aspect ratio of the nuclei during transport (inside the tube) and after being expelled (outside the tube), which represents the extent of nuclear deformation during the process. Error bars represent standard deviation (n=7), ****p<0.0001 (paired Student’s t-test).

Next, we shifted our attention to monitoring the nucleus as it moves through the PT. Given the discrepancy in diameter between the nucleus and the PT, we hypothesized that either 1) the PT is flexible enough that it could deform to accommodate the nuclei, or 2) the nuclei must deform to fit the PT diameter (**Fig. 5b**). To assess which hypothesis may be correct, we performed live-cell imaging of *A. algerae* PT germination *in vitro*, and monitored the nuclei as they traversed through the PT. Our results show gross deformation of the nuclei on a millisecond timescale as they travel through the PT (**Fig. 5d, Supplementary Fig. 10b, Supplementary Video 11**). However, after exiting the tube, the nuclei return to a globular shape (**Fig. 5d**). To better quantify this nuclear deformation, we calculated the aspect ratio (the shortest dimension divided by the longest dimension) of the nuclei inside the PT and after exit from the PT (**Fig. 5e**). The aspect ratio should be close to 1 for an isotropic object (e.g. circle), whereas an aspect ratio less than 1 indicates an elongated object (ellipse). The average aspect ratio of the paired nuclei is 0.11 ± 0.04 inside the tube and 0.87 ± 0.07 after they are expelled, outside the PT (**Fig. 5e**). The nucleus travels through the tube at remarkable velocities, approaching 270 ± 115 μm.s^-1^ (**Supplementary Fig. 10d**), which is comparable to PT firing velocities in *A. algerae*. Of note, the nucleus pauses within the PT during translocation in all seven movies we recorded *in vitro* (**Fig. 5d, Supplementary Fig. 10b,c, Supplementary Video 11**). This may reflect a change in the forces that are driving cargo transport through the PT, though it remains to be determined whether pausing also occurs during germination *in vivo*.

## Discussion

### Mechanistic insights into polar tube germination

Microsporidia comprise an entire phylum of extremely successful parasites, all of which share the common feature of the polar tube infection apparatus. Our data combine spatial and temporal information to provide new insights about how the PT is packed in dormant spores, the PT germination process, and synchronization of PT firing and cargo transport. Synthesizing these data, we present a model for PT germination (**Fig. 6**), in which we can clearly define some aspects of the germination process, while others still remain ambiguous. We focus on *A. algerae*, as we have the most information for this species. In the dormant spore, the PT is a right-handed helix packed at an angle relative to the A-P axis of the spore, and interacts closely with the vacuole. The two nuclei in *A. algerae* are nestled in the bowl-shaped vacuole, and surrounded on the sides by the coiled PT (**Fig. 6a**). Previous data suggest that triggering germination is dependent on osmotic pressure buildup within the spore^56^, which may initiate the rupture of the spore wall and PT firing^57^. Once triggered, the PT fires and reaches its full length in under 1 second (T_EXT90_ = ∼830 ms). By tracking the nuclei during this germination process, we find that as the germination process initiates, the two nuclei begin to rearrange in the spore (**Fig. 6b**, step 3). Approximately 500 ms after PT firing is initiated, the nuclei exit the spore body together, and deform drastically to fit into the PT. The average time to 50% extension (T_EXT50_) in *A. algerae* is approximately 600 ms, suggesting that once the tube has reached ∼50% extension, cargo transport is initiated. This observation is consistent with a model in which the PT everts^17,58^, which would only allow cargo to enter the tube after 50% extension. The velocity at which the cargo is transported through the tube is comparable to the velocity of PT extension, and the cargo regains its initial shape after exiting the PT. Many open questions remain and further studies will be required to definitively address the mechanistic basis of PT germination.

**Fig. 6.**
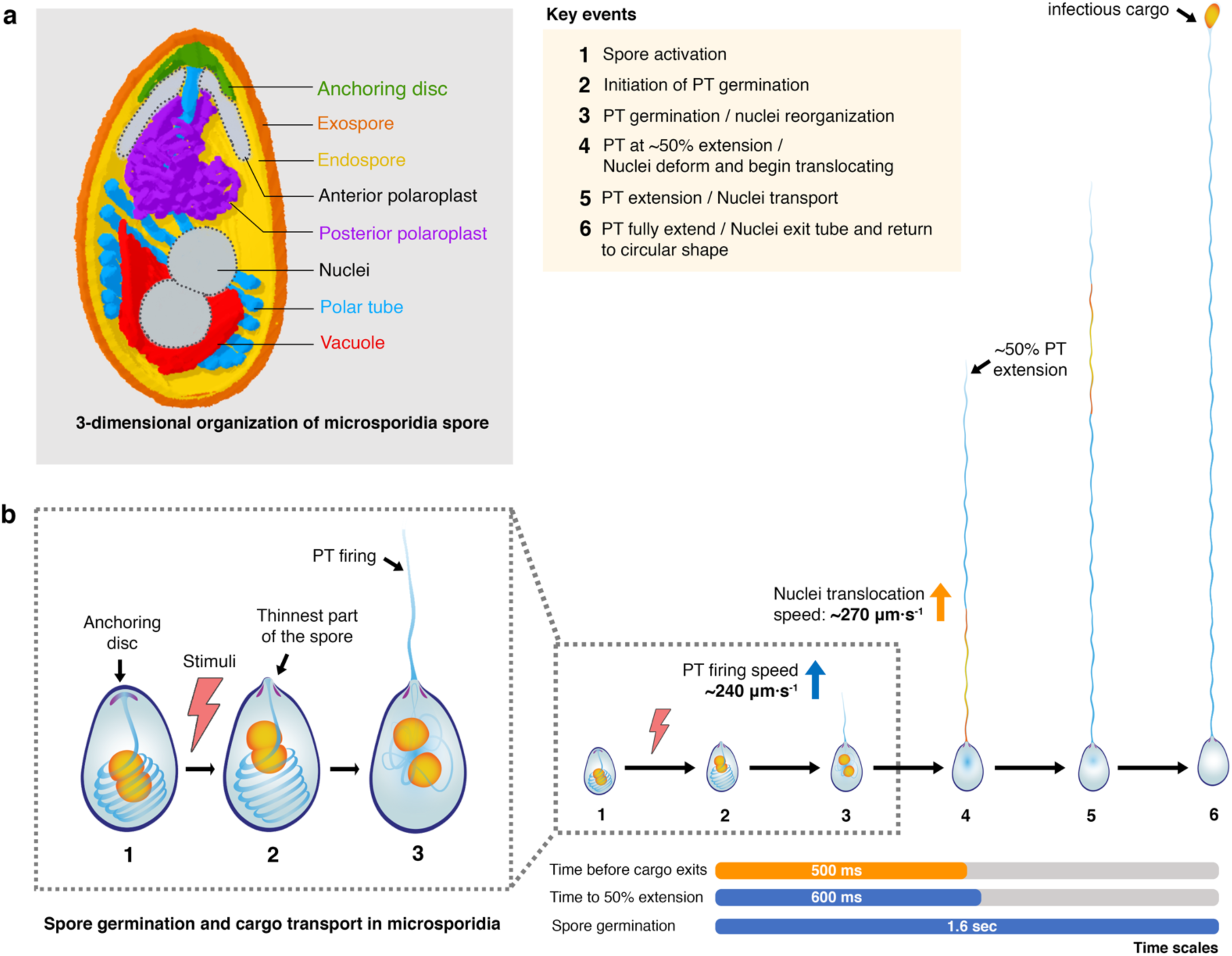
Model for PT germination and nuclear transport through the PT. **(a)** Model for 3-dimensional organization of a microsporidian spore. Combining our SBFSEM and 2D TEM data for both *A. algerae* and *E. hellem*, we generate a composite 3D model that shows spatial organization of organelles inside the microsporidian spore. Elements that were not observed in the *A. algerae* SBFSEM data (anterior polaroplast and nuclei) have been added manually and are in gray. The anterior part of the PT (blue) is straight and connected to the anchoring disc (green). This part of the PT is surrounded by the anterior (gray) and posterior (purple) polaroplast. The posterior part of the PT is coiled and packed at an angle relative to the A-P axis. The PT resembles a rib cage around the vacuole (red) and nuclei (gray). The vacuole is bowl shaped, and localized beneath the nuclei. **(b)** Model for PT germination and cargo transport in *A. algerae*. (1) The spore is triggered in the presence of a stimulus (*in vivo* triggers remain largely unknown; *in vitro* triggers are described in **Methods**). (2) PT firing is initiated at the thinnest part of the spore coat, where the anchoring disc is localized. (3) After initiation of PT firing and during Phase 1 (PT elongation), the nuclei, and presumably other organelles, are reoriented. (4) At the stage in which the PT is extended to just past 50%, the nuclei deform to fit into the PT and exit the spore. (5) Nuclei (and likely other cargo) are translocated through the PT, at a speed comparable to that of PT extension. (6) The nuclei exit the PT and regain a circular shape at the tip of the PT.

### Rapid nuclear deformation in microsporidia

In our study, we observed incredibly fast and large deformation of the nuclei as they traveled through the PT. This is reminiscent of other cell translocation and migration processes, such as immune cells squeezing through tight junctions to exit the bloodstream, or tumor cells penetrating tissues^59–61^. Nuclear deformation is necessary to facilitate tumor cell invasion, as the cells must adapt their shape to accommodate available space within the three-dimensional architecture of the tissue. Two key differences between the nuclear deformation we observe in microsporidia and analogous processes in tumor cell invasion is that the timescale is much faster in microsporidian nuclear transport (milliseconds compared with minutes/hours), and the microsporidian nuclear distortion is much more severe, as assessed by the aspect ratio. During tumor cell invasion and nuclear deformation, DNA damage is reported to occur^62,63^, and it therefore seems plausible that the microsporidian genome may also be subject to shearing and other stresses.

If extensive damage to the microsporidian genome does occur, efficient repair mechanisms must be in place to allow the rapid activation of transcription and DNA replication upon cargo delivery into the host cell. In migrating cancer cells, nuclear envelope breakdown has also been noted^62,63^, which likely enhances deformability of the packaged DNA. Some cancer cells also express lower levels of lamins^64–67^, which are known to be important cytoskeletal proteins that rigidify the nuclear envelope. While cells expressing low levels of lamin A show a high migration rate through tissues^68^, overexpression of lamin A has been shown to result in reduced nuclear deformability and impaired cell passage through narrow constrictions^69^, consistent with the idea that the presence of lamins contributes substantially to rigidity of the nucleus. In agreement with previous reports that lamins are only present in Metazoans^70,71^, we were unable to identify any lamins or homologs in fully sequenced microsporidian genomes^13,72,73^. This suggests that microsporidia may lack lamins entirely, perhaps explaining why nuclear deformation occurs to a much higher degree than in mammalian cells, and on a rapid timescale. Future studies will reveal the nature of cargo transport and address questions of the chromatin state during nuclear transport, whether the nuclear envelope is intact, and how cargo is modulated once delivered to the host.

## Methods

### Propagation of microsporidian parasites

To propagate the parasites, microsporidia *A. algerae* (ATCC^®^ PRA-168™), *E. hellem* (ATCC^®^ 50504™), and *E. intestinalis* (ATCC^®^ 50506™) were grown in Vero cells (ATCC^®^ CCL-81™) using Eagle’s Minimum Essential Medium (EMEM) (ATCC® 30-2003™) with 10% heat-inactivated fetal bovine serum (FBS) at 37°C and with 5% CO_2_. At 70%-80% confluence, parasites were added into a 25 cm^2^ tissue culture flask and the media was switched to EMEM supplemented with 3% FBS. Cells were allowed to grow for fourteen days and medium was changed every two days. To purify spores, the infected cells were detached from tissue culture flasks using a cell scraper and placed into a 15 ml conical tube, followed by centrifugation at 1,300 g for 10 min at 25°C. Cells were resuspended in sterile distilled water and mechanically disrupted using a G-27 needle. The released spores were purified using a Percoll gradient^73^. Equal volumes (5 mL) of spore suspension and 100% Percoll were added to a 15 mL conical tube, vortexed and then centrifuged at 1,800 g for 30 min at room temperature. The purified spore pellets were washed three times with sterile distilled water and stored at 4°C in distilled water for further analyses.

### Germination conditions for microsporidia spores

Germination conditions, to activate PT firing, are different among microsporidia species^46,47,74^. For *A. algerae*, the germination buffer was slightly modified from the previously reported condition^46^. Spore germination was triggered using 10 mM Glycine-NaOH buffer pH 9.0 and 100 mM KCl, yielding ∼70% germination. For both *Encephalitozoon* species, spore germination was triggered using germination buffer containing 140 mM NaCl, 5 mM KCl, 1 mM CaCl_2_, 1 mM MgCl_2_, and 5% (v/v) H_2_O_2_ at pH 9.5, as previously described^47^. Germination buffer was freshly prepared for each experiment.

### Electron microscopy

Purified spores were fixed in 0.1 M sodium cacodylate buffer (pH 7.2) containing 2.5% glutaraldehyde and 2% paraformaldehyde. For transmission electron microscopy, fixed spores were post-fixed with 2% osmium tetroxide (OsO_4_), embedded in 2% agar, block stained in 1% aqueous uranyl acetate, and dehydrated using a gradient of cold ethanol. The samples were then transferred in propylene oxide and embedded in EMbed 812 (Electron Microscopy Sciences, catalog #14121). 70 nm thin sections were cut, mounted on copper grids and stained with uranyl acetate and lead citrate. Stained grids were inspected using a Thermo Fisher Talos 120C electron microscope and imaged with a Gatan OneView camera using a nominal magnification of 22,000x corresponding to a pixel size of 0.652 nm/pixel on the specimen.

For serial block-face scanning electron microscopy (SBFSEM), the sample block was mounted on an aluminum 3View pin and electrically grounded using silver conductive epoxy (Ted Pella, catalog #16014). The entire surface of the specimen was then sputter coated with a thin layer of gold/palladium and imaged using the Gatan OnPoint BSE detector in a Zeiss Gemini 300 VP FESEM equipped with a Gatan 3View automatic microtome. The system was set to cut 50 nm slices, imaged with gas injection setting at 40% (2.9 × 10^−3^ mBar) with Focus Charge Compensation to reduce electron accumulation charging artifacts. Images were recorded after each round of sectioning from the blockface using the SEM beam at 1.5 keV with a beam aperture size of 30 µm and a dwell time of 1.0 µs/pixel. Each frame is 22 × 22 µm with a pixel size of 2.2 nm. Data acquisition occurred automatically using Gatan DigitalMicrograph (version 3.31) software. A stack of 150 slices was aligned and assembled using Fiji^75^. A total volume of 22 × 22 × 11 µm^3^ was obtained from the sample block.

Segmentation of organelles of interest, 3D reconstruction, and quantification of the spore size, volumes and PT length in the intact spores were performed using Dragonfly 4.1 software (Object Research Systems, ORS). SBFSEM sections were automatically aligned using SSD (sum of squared differences) method prior to segmentation. Graphic representation of the spores and PT were performed with either Dragonfly software or UCSF Chimera^76^.

Measurement of the PT angles and the distance between coils were made in UCSF Chimera^76^. To facilitate this, two atoms were placed as markers on the anterior-posterior ends of the spore to create an A-P axis. 3 more atoms were placed along the PT coil to generate a plane corresponding to the PT. Then, the PT angles were measured between the A-P axis and the PT plane. To measure the distance between the coils, atoms were manually placed at the center of two adjacent coils, and the distance between these was measured.

### Optical microscopy

#### Live-cell imaging of PT firing

2 μl of purified spores (∼10^8^ spores/ml) were mixed with 10 ul of germination buffer. The reaction was placed on ice to prevent PT firing prior to imaging. 2 ul was placed on a poly-L-lysine-coated glass slide (Fisher Scientific, catalog #12-545-78) and sealed with a #1.5 18 × 18 mm coverslip (Fisher Scientific, catalog #12-519-21A). Polar tube firing typically occurred ∼2-5 minutes after mixing the spores with the germination buffer. PT firing was imaged using a Nikon Eclipse Ti microscope with a Nikon 60x N.A. 1.4 oil immersion Plan Apochromat Ph3 phase-contrast objective lens. An Andor Zyla 5.5 megapixel sCMOS camera was used, which provided a wide field of view at 14-50 frames per second with 3-35 ms exposure time, no binning was applied. The microscope was equipped with an environmental chamber which was set at 30°C for these experiments.

#### Fixed cell imaging

4 μl of *A. algerea* spores (10^8^ spores/ml) was incubated with 20 μl of germination buffer at 30°C for 30 min. 40 ul of NucBlue™ Live ReadyProbes™ Reagent (Invitrogen, catalog #R37605) was added and the reaction was incubated at 25°C for 20 min. Spores were pelleted by centrifugation at 1,000 g for 1 min at room temperature and the supernatant was removed. Spores were resuspended in 6 ul of fresh germination buffer. 2 μl of the reaction was placed onto a glass slide and sealed with a #1.5 18 × 18 mm coverglass. Spores were imaged using a Nikon Eclipse Ti microscope with a Nikon 60x N.A. 1.4 oil immersion Plan Apochromat Ph3 phase-contrast objective lens. A Zyla 5.5 megapixel sCMOS camera was used at 126 ms exposure time, and no binning was applied.

#### Live-cell imaging of nuclear transport

4 μl of purified spores (6×10^7^ spores/ml) were mixed with 40 μl of NucBlue (Invitrogen, catalog #R37605) and incubated at 25°C for 40 min. Spores were then centrifuged at 5,000 g for 1 min at room temperature and the supernatant was removed. 10 μl of germination buffer was added to the pre-stained spores and stored on ice prior to imaging. 2 ul of this reaction was added to a poly-L-lysine coated glass slide and covered with a #1.5 18 × 18 mm coverslip.

To image nuclear movement inside the spore coat prior to translocation into the PT, imaging was performed on a Nikon Eclipse Ti microscope with Nikon 60x N.A. 1.4 oil immersion Plan Apochromat Ph3 objective lens. Intensity of fluorescent excitation and intensity of transmitted light were balanced to allow simultaneous single channel single camera imaging (Duo-detection). A Zyla 5.5 megapixel sCMOS camera was used, providing a wide field of view at 28 frames per second with 30 ms exposure time, no binning was applied.

To observe nuclear translocation through the PT, imaging was performed on a Zeiss AxioObserver Z1 with 40x N.A. 1.3 EC Plan-Neofluar oil immersion objective lens. An Axiocam 503 Monochrome CCD camera was used, yielding 20 frames per second with 45 ms exposure time, and 3×3 binning was applied.

#### Image analysis

Kymographs of the PT germination were generated from movies using Fiji software^75^ with the straighten function. Measurement of the PT length was quantified from raw time-lapse images using Fiji software^75^. The PT was traced using the segmented line function. The maximum PT length was defined from the exit site from the spore body to the point where the infectious cargo connects to the PT. Velocity and acceleration of the PT firing process were calculated by Δy/Δx, where Δy is changes in PT length or changes in velocities, and Δx is the corresponding time interval. Graphs were plotted using GraphPad Prism 8 software. Fitting of data in **Fig. 4b** was performed using a ‘sigmoidal, 4PL, X is concentration’ equation. R^2^ values were 0.89 for *A. algerae*, 0.60 for *E. hellem*, and 0.93 for *E. intestinalis*.

For nuclear translocation, the kymographs were analyzed using Fiji software^75^. Speed of nuclear translocation was measured from changes of the distance of nuclear signals divided by the corresponding time interval. The aspect ratio of the nuclear deformation was quantified by the width (W) divided by the height (H) of nuclear signals from seven kymographs. The aspect ratio for the nucleus in the PT is measured during the pause phase of transport to minimize the impact of blurring due to movement, and the aspect ratio for the nucleus outside the tube is measured when it emerges at the distal end of the PT.

### Validation of PT length comparison between pre-germination and post-germination states

Our measurements of PT length inside the spore (pre-germination) are made from analyzing SBFSEM data, while measurements of the PT outside the spore (post-germination) are made using optical microscopy. To assess whether measurements from these methods are comparable, we measured the spore length for both *A. algerae* and *E. hellem* from both these techniques. The spore length obtained from SBFSEM and LM are similar in both species (3.8 ± 0.4 μm from SBFSEM and 3.7 ± 0.4 μm from LM for *A. algerae*; 2.8 ± 0.3 μm from both SBFSEM and LM for *E. hellem*). These results validate comparing measurements between the two methods.

### Statistical analyses

GraphPad Prism 8 software was used for all statistical analyses. In all analyses, we used a two-tailed unpaired Student’s t-test to compare the difference between two groups, with the exception of two analyses: 1) PT shortening after germination and 2) the nuclear aspect ratio. For these two exceptions a two-tailed paired Student’s t-test was used. P values are reported in the figure legends.

## Acknowledgements

We thank James J. Becnel and Neil Sanscrainte for sharing initial samples and expertise for propagation of *A. algerae* spores; Kristen Dancel-Manning for helping with figure preparation; Chris Petzold at the NYU Microscopy Core for assistance with preparation of EM samples; Huilin Li from the NYU Biostatistics Resource for guidance with statistical analysis and Nicolas Coudray for assistance with analysis in Chimera. We thank Emily Troemel, Louis Weiss, Alex Mogilner, Saima Sidik, Georgia Isom, Juliana Ilmain, Noelle Antao and Frederick Rubino for critical reading and feedback on our manuscript, and all members of the Bhabha/Ekiert labs as well as Manu Prakash for helpful discussions. We gratefully acknowledge the following funding sources: 19POST34430065 (American Heart Association, to P.J.), R35GM128777 (NIGMS, to D.C.E.), PEW-00033055 (Pew Biomedical Scholars, to G.B.), SSP-2018-2737 (Searle Scholars Program, to G.B.), R01AI147131 (NIAID, to G.B.). The NYU Microscopy Core is partially supported by NYU Cancer Center Support Grant NIH/NCI P30CA016087, and Zeiss Gemini 300 SEM with 3View was purchased with support of NIH S10 ODO019974-01A1.

## Data availability

All live-cell imaging data have been deposited on an open access digital library in Zenodo,DOI:10.5281/zenodo.3707829

## Competing interests

The authors declare that they have no conflict of interest.

## Contributions

P.J., F.L., D.C.E., and G.B. conceived the study. P.J., M.C., J.S., and M.U. carried out the experiments. P.J., A.D., M.U., G.B. and D.C.E. analyzed the data. P.J., D.C.E., and G.B. wrote the manuscript with contributions from all other authors.

## Supplementary figures

**Supplementary Figure 1.**
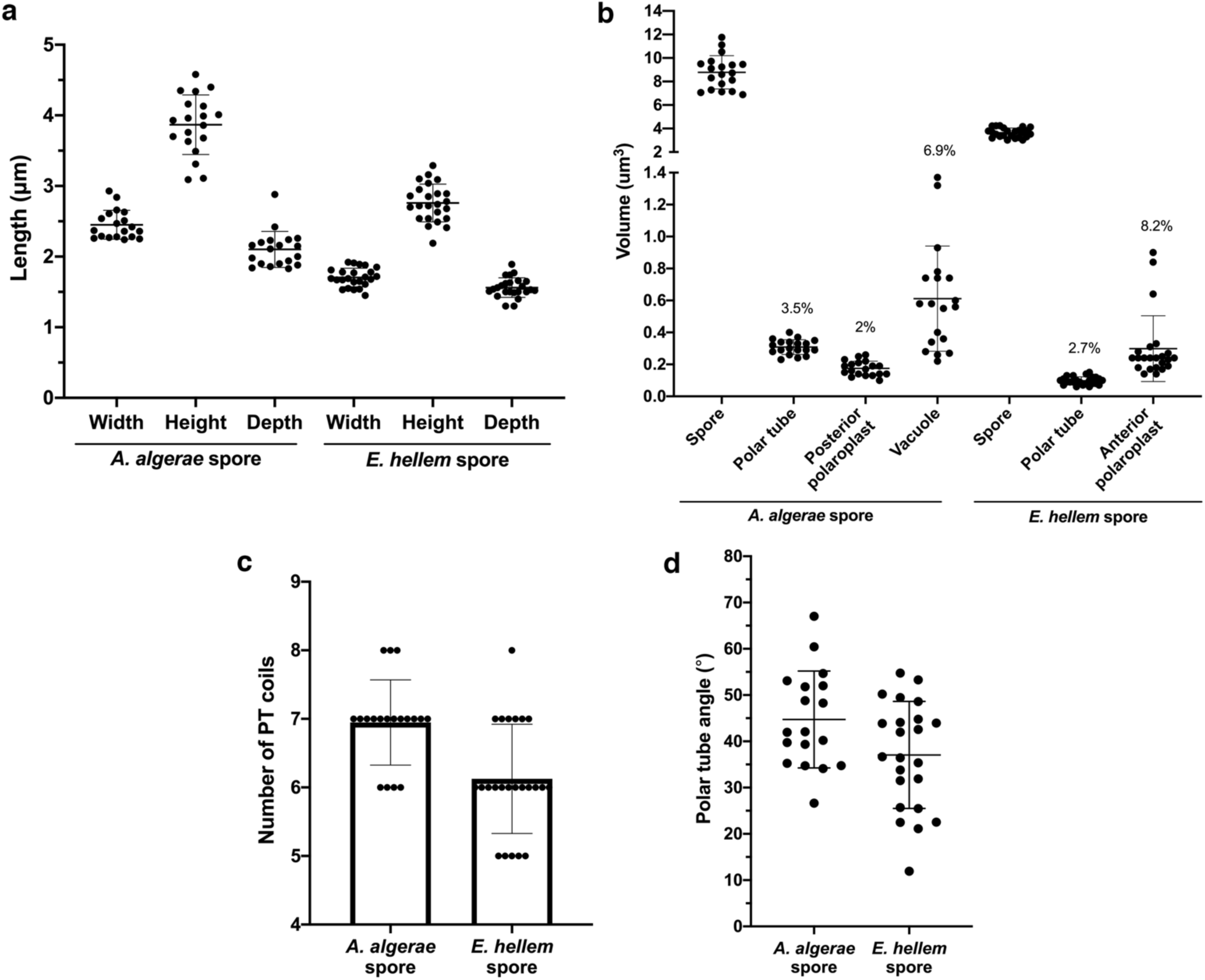
Quantification of volume and dimensions of spore organelles from SBFSEM. **(a)** Quantification of spore dimensions. **(b)** Quantification of volumes. Organelle volumes as a percentage of the entire spore volume are noted on the graph **(c)** Number of PT coils quantified from 3D reconstruction of spores. **(d)** Quantification of the angle between PT coils and the A-P axis. All error bars in this figure represent standard deviation (n=19 for *A. algerae* and n=23 for *E. hellem*).

**Supplementary Figure 2.**
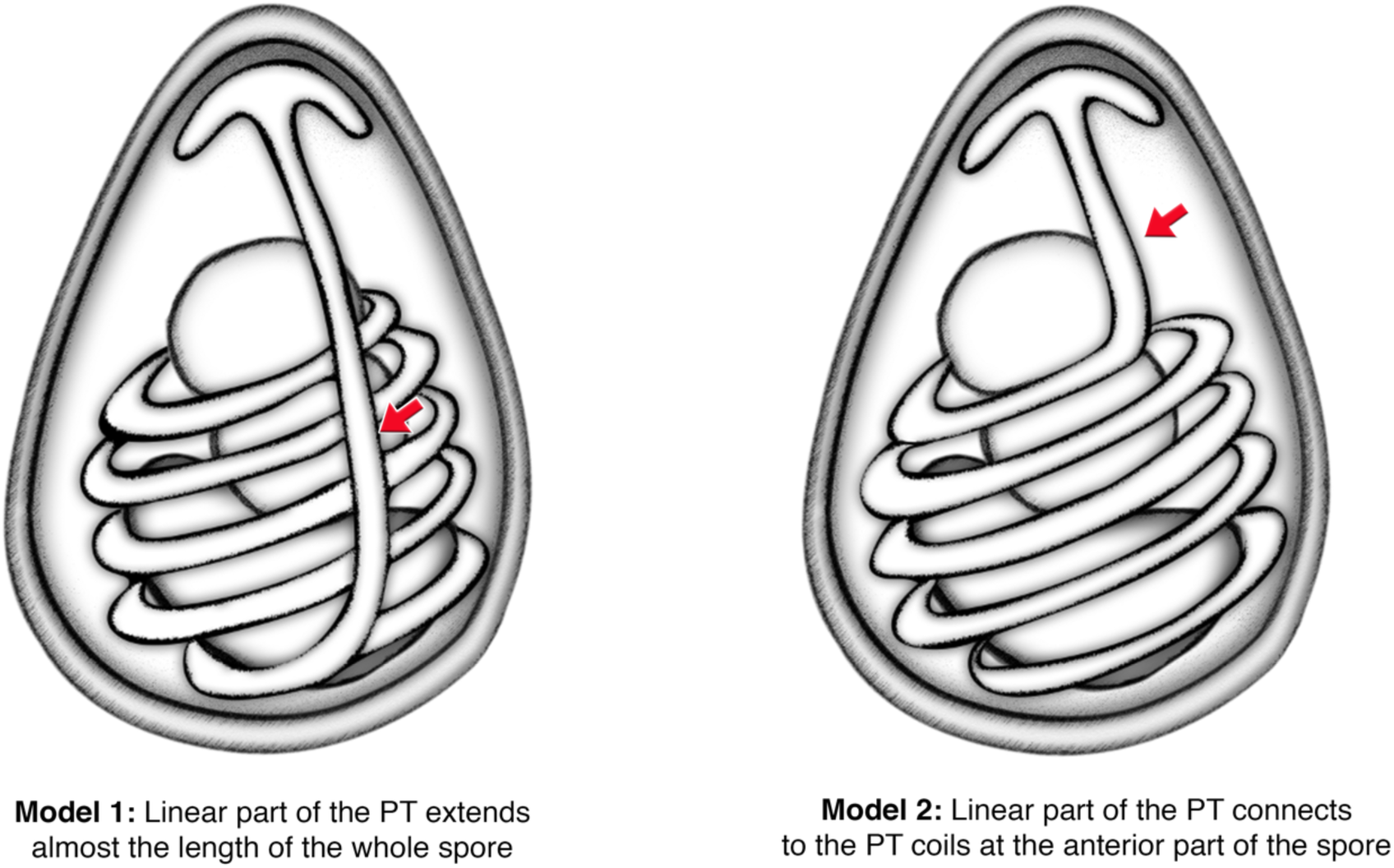
Possible models of how the linear and coiled parts of the PT are connected. Schematic diagrams showing two possible models of the connection between straight and coiled regions of the PT. Red arrows indicate the region where the PT is straight. Model 1 is adapted from Cali *et al*^*39*^.

**Supplementary Figure 3.**
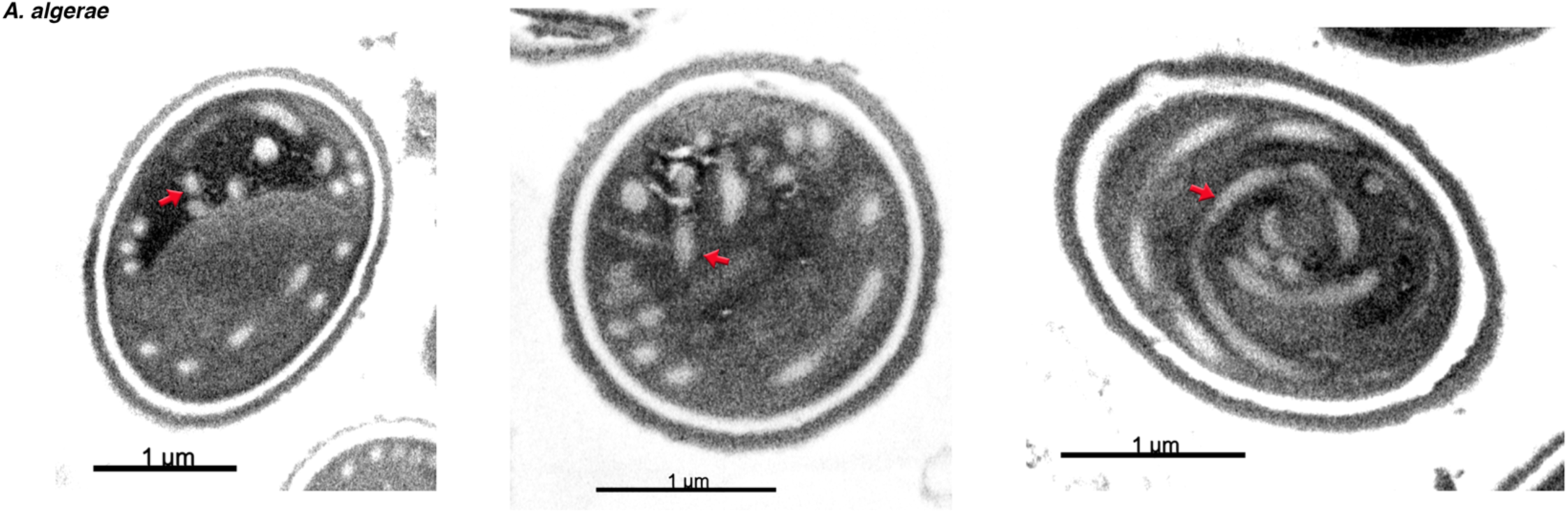
Representative SBFSEM slices through spores with tangled PT ends. Three representative SBFSEM sections originating from spores with tangled PT ends, as described in Fig. 2d. Red arrows indicate the PT.

**Supplementary Figure 4.**
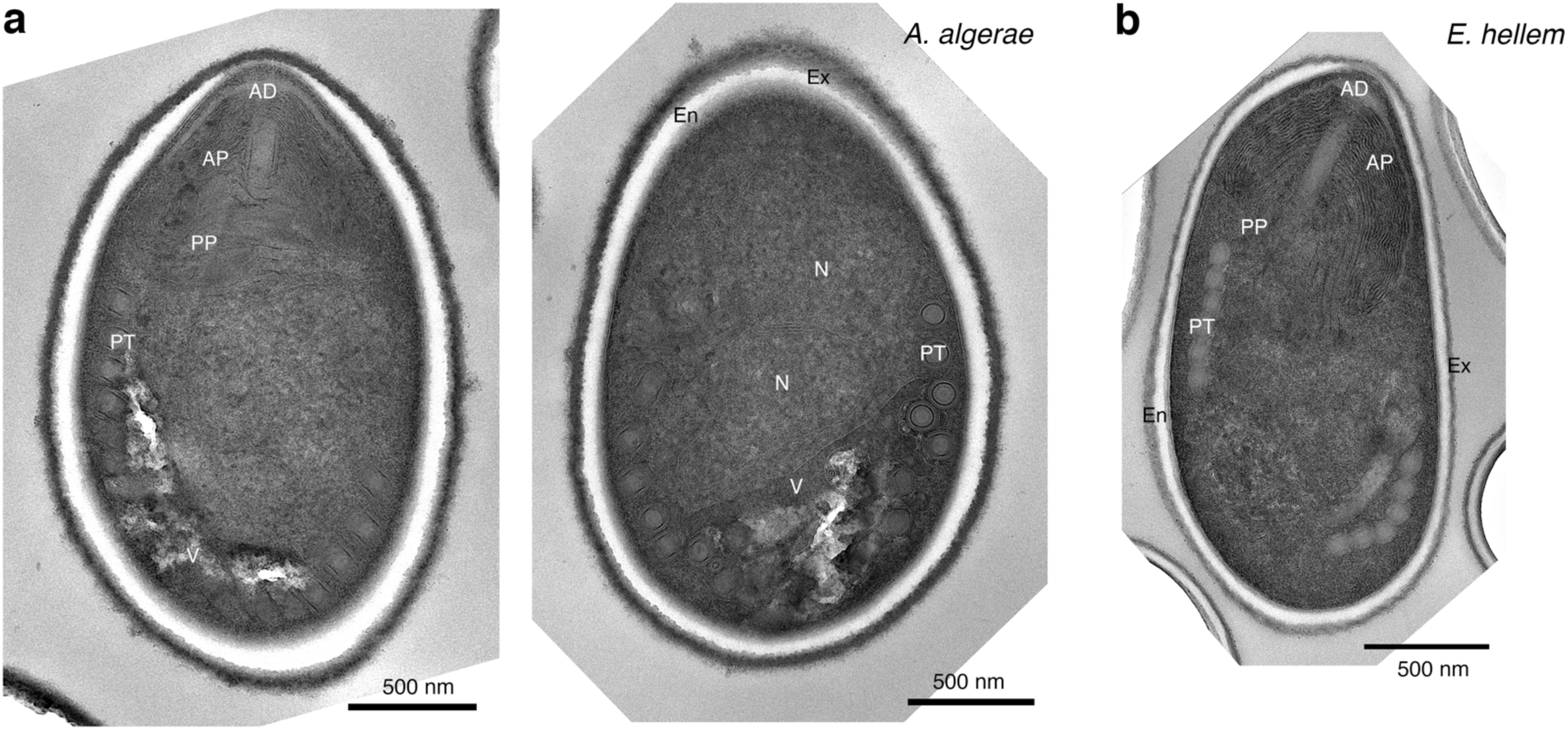
2D TEM images of *A. algerae* and *E. hellem* spores. TEM sections of **(a)** *A. algerae* spores and **(b)** *E. hellem* spore indicating structures inside the spore, including exospore (Ex), endospore (En), anchoring disc (AD), anterior polaroplast (AP), posterior polaroplast (PP), polar tube (PT), nucleus (N), and vacuole (V). These samples were used for SBFSEM experiments. The right panel of (a) is the same as that shown in Fig. 2e, but without color overlay.

**Supplementary Figure 5.**
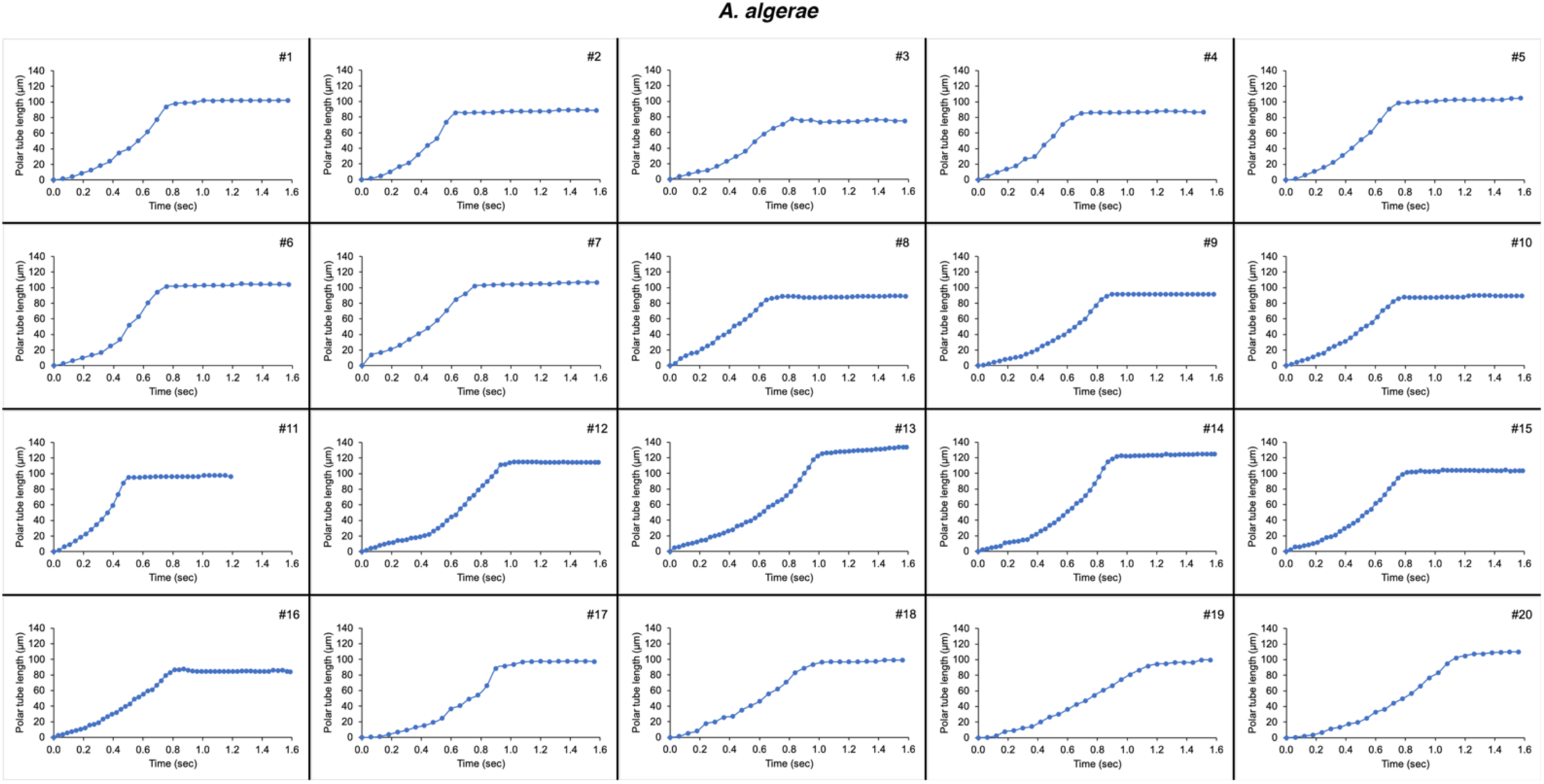
*A. algerae* PT length as a function of time during the germination process. Graphs represent polar tube length over the time period of PT germination for 20 individual spores. See Supplementary data file 1 for data used to generate these plots.

**Supplementary Figure 6.**
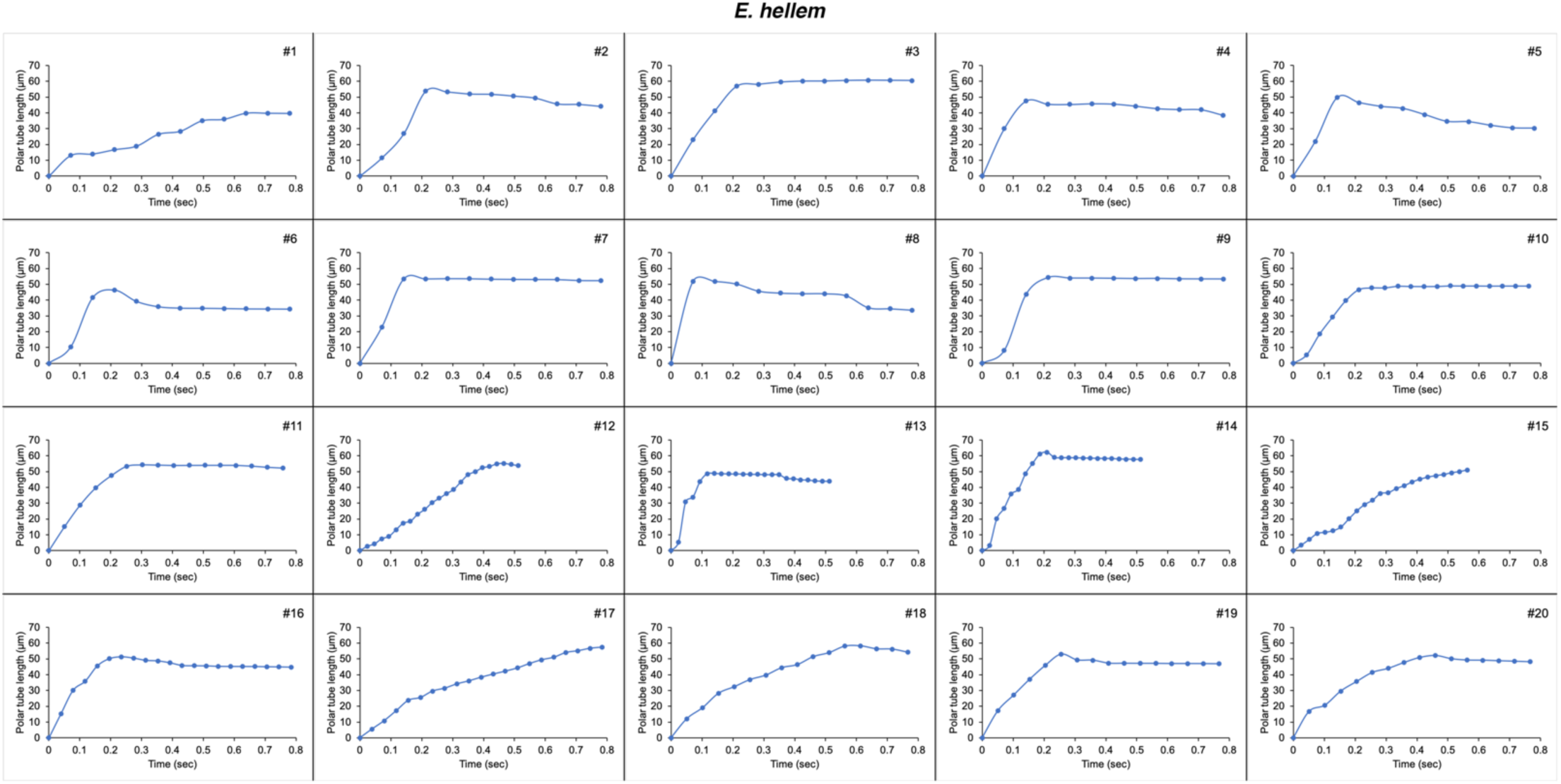
*E. hellem* PT length as a function of time during the germination process. Graphs represent polar tube length over the time period of PT germination for 20 individual spores. See Supplementary data file 1 for data used to generate these plots.

**Supplementary Figure 7.**
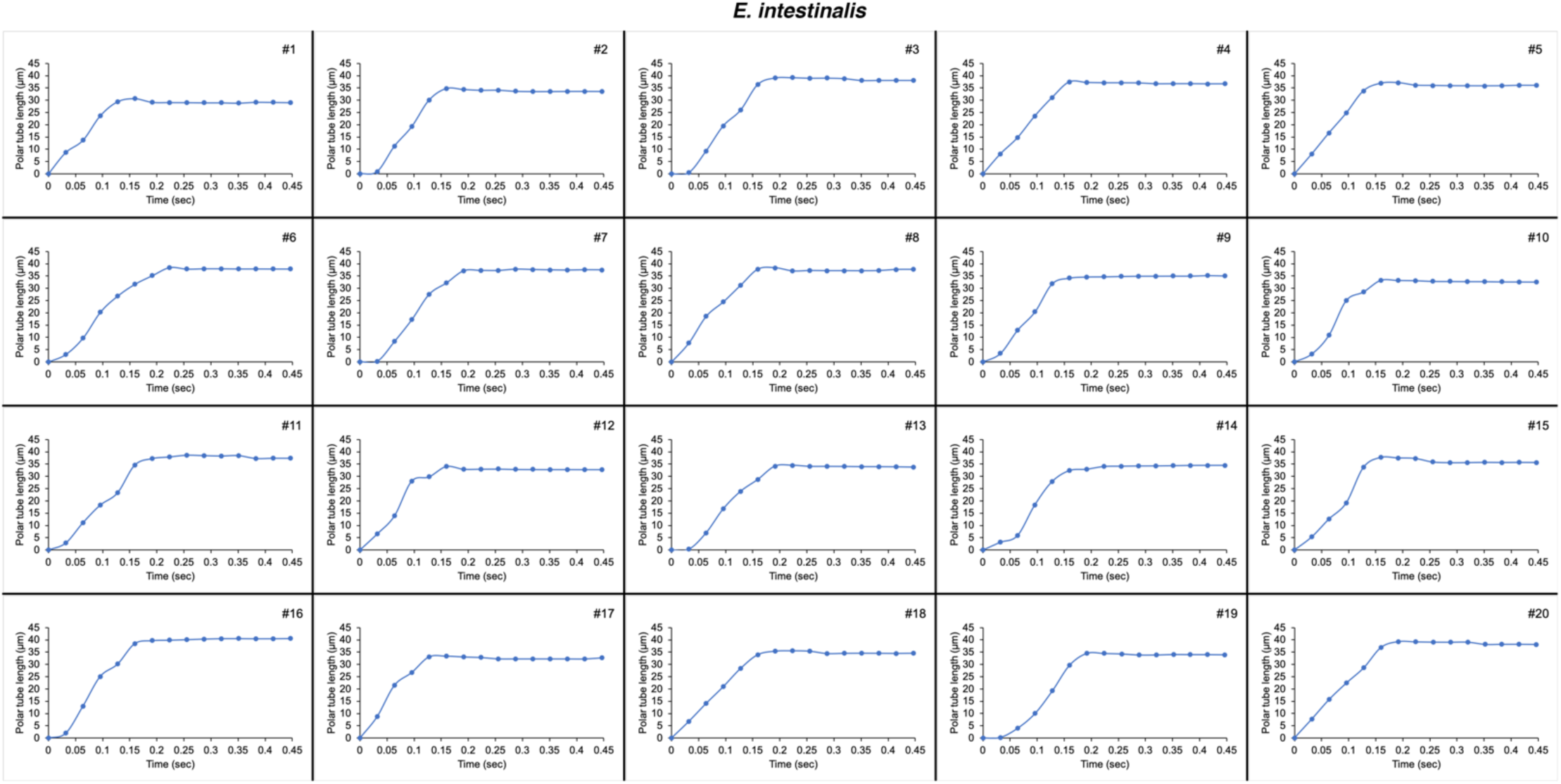
*E. intestinalis* PT length as a function of time during the germination process. Graphs represent polar tube length over a period of PT germination from 20 individual spores.

**Supplementary Figure 8.**
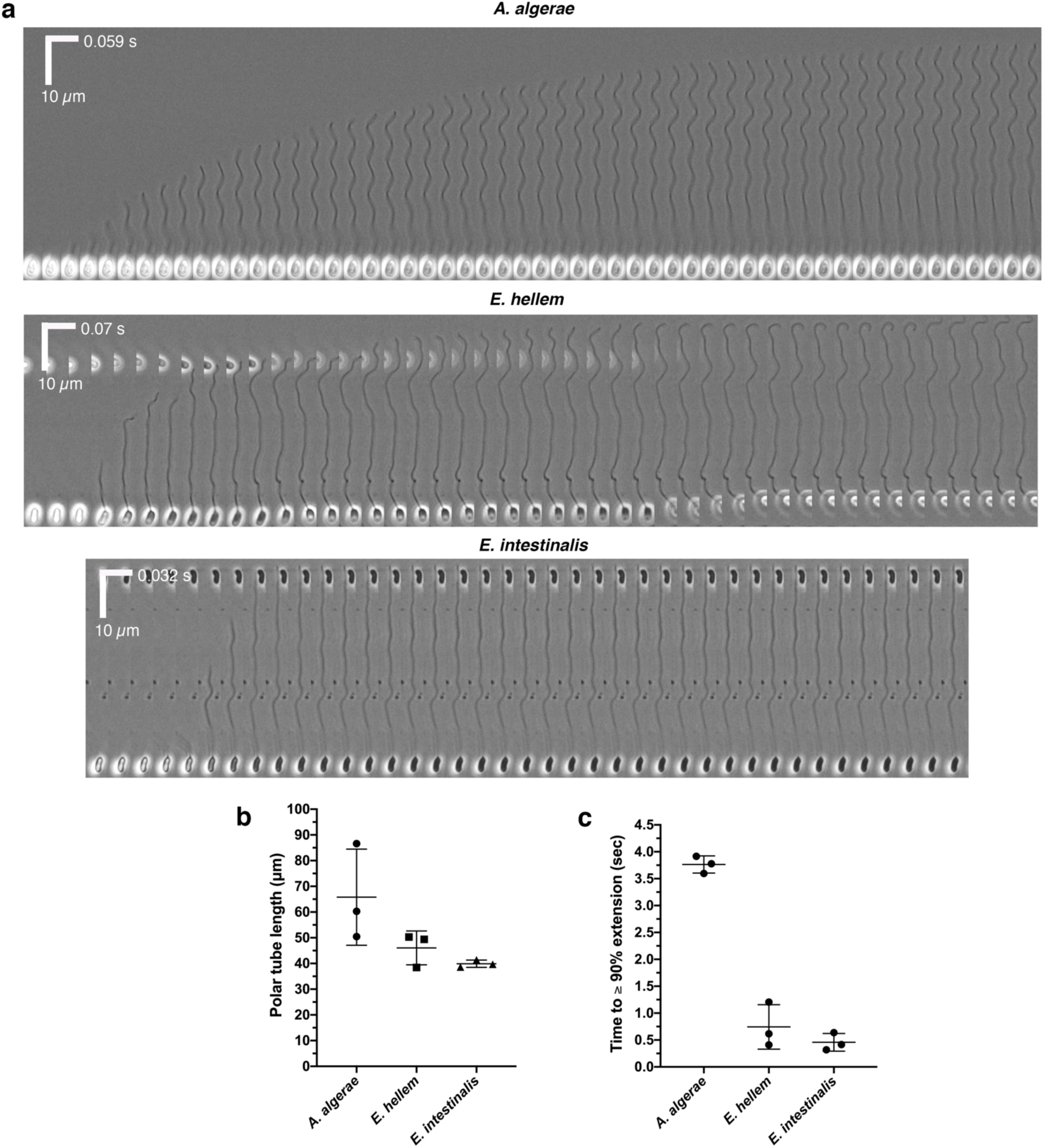
Examples of incomplete germination events. **(a)** Kymographs of incomplete spore germination from *A. algerae, E. hellem*, and *E. intestinalis*. Scale bar for time is shown on the X-axis and for distance on the Y-axis. For incomplete germination events observed, the distal end of the tube appeared to be straight instead, in contrast to the hooked ends usually observed for complete germination^54^. **(b)** PT length quantified from incomplete germination events. **(C)** Quantification of the time for the PT to reach ≥ 90% of its maximum length. The error bars in this figure represent standard deviation (n=3 for each species).

**Supplementary Figure 9.**
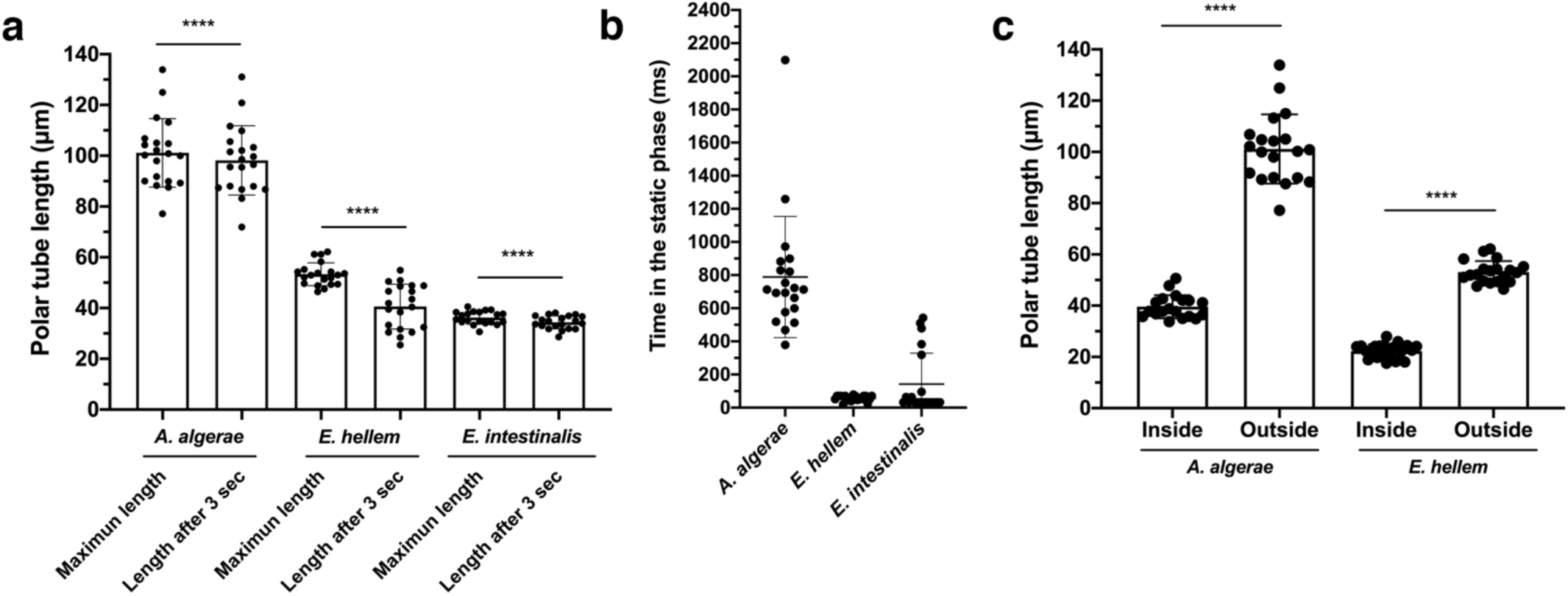
Quantification of PT lengths at different stages, and time spent in phase 2 of the PT germination. **(a)** Graph showing the maximum length and the length of the PT at 3 sec after germination is complete. The data in this graph are the same as in Fig. 4f, but not normalized. ****p<0.0001 (paired Student’s t-test). **(b)** Time spent in phase 2 of germination, when the PT is static. **(c)** Quantification of PT length in the intact spores from SBFSEM and the length of fully extended PT from optical microscopy. The graph is the same as presented in Fig. 4g, but with individual data points plotted. ****p<0.0001 (unpaired Student’s t-test). Error bars in this figure represent standard deviations (n=20 for each species).

**Supplementary Figure 10.**
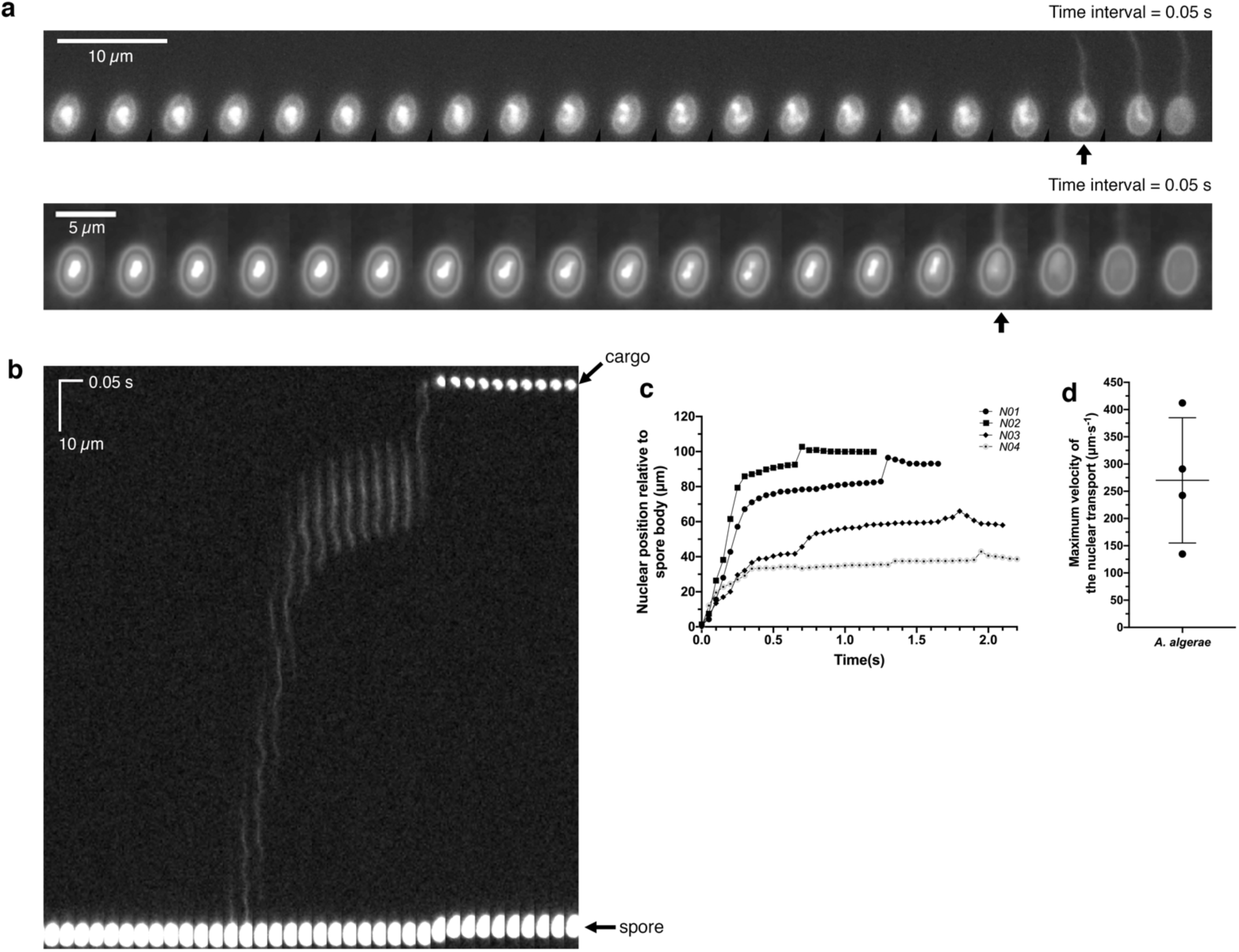
Nuclei imaged during PT germination. **(a)** Additional time-lapse images of fluorescently labeled nuclei inside spores during PT germination. Top and bottom represent two individual spores. Time intervals are 50 ms. Black arrows indicate the frame in which the nuclei have begun to enter the PT. **(b)** Additional kymograph of the nuclear translocation through the PT. **(c)** Quantification of the nuclear position relative to spore coat over time (n=4). **(d)** Graph showing maximum velocity of the cargo transport process. The data was calculated from (c). Error bar represents standard deviation (n=4). A total of seven movies were recorded to monitor nuclear transport. However, the nuclear signal in the first few frames of the movie was below our detection threshold for three of these movies, which have not been quantified here.

## Supplementary videos

**Supplementary Video 1. 3D reconstruction of A. *algerae* spore from SBFSEM**. Representative 3D reconstruction of an *A. algerae* spore. At the beginning of the Video, slices through the spore are shown. Each color represents an individual organelle: exospore (orange), endospore (yellow), PT (blue), vacuole (red), posterior polaroplast (purple), and anchoring disc (green).

**Supplementary Video 2. 3D reconstruction of *E. hellem* spore from SBFSEM**. Representative 3D reconstruction of an *E. hellem* spore. Each color represents an individual organelle: exospore (orange), endospore (yellow), PT (blue), anterior polaroplast (pink), and anchoring disc (green).

**Supplementary Video 3. Live-cell imaging of *A. algerae* PT germination**. Time-lapse video of PT germination in *A. algerae*, corresponding to Fig. 4a.

**Supplementary Video 4. Live-cell imaging of *E. hellem* PT germination**. Time-lapse video of PT germination in *E. hellem*, corresponding to Fig. 4a.

**Supplementary Video 5. Live-cell imaging of *E. intestinalis* PT germination**. Time-lapse video of PT germination in *E. intestinalis*, corresponding to Fig. 4a.

**Supplementary Video 6. Live-cell imaging of *E. hellem* PT shortening after germination**. Time-lapse video of the PT germination in *E. hellem*, corresponding to Fig. 4f. After the emergence of the infectious cargo at the end of the PT, the PT starts to shrink. However, rather than a synchronized shortening of the entire tube across its complete length, segments of the tube appear to contract at slightly different times.

**Supplementary Video 7. Incomplete germination event for *A. algerae***. Time-lapse video of incomplete PT germination in *A. algerae*.

**Supplementary Video 8. Incomplete germination event for *E. hellem***. Time-lapse video of incomplete PT germination in *E. hellem*.

**Supplementary Video 9. Incomplete germination for *E. intestinalis***. Time-lapse video of incomplete PT germination in *E. intestinalis*.

**Supplementary Video 10. Nucleus movement inside the spore prior to translocation through the PT**. Time-lapse images of *A. algerae* nucleus inside the spore. Dual detection of NucBlue fluorescence and transmitted light was applied in order to visualize both the DNA and PT simultaneously.

**Supplementary Video 11. Nucleus movement through the PT**. Time-lapse video of *A. algerae* nuclear transport through the PT.

## Notes

DOI:10.5281/zenodo.3707829

